# Filling a metabolism data gap in marine fish reveals dual pace-of-life and reproductive strategy axes Fish Metabolism and Pace-of-Life

**DOI:** 10.64898/2026.09.28.755002

**Authors:** Marine Beneat, Alaia Morell, Fabien Moullec, Nicolas Barrier, Yunne-Jai Shin, Bruno Ernande

## Abstract

Resting metabolic rate, the basal energy required for an organism’s maintenance, is central to predicting the eco-evolutionary consequences of environmental and anthropogenic pressures on marine ectotherms. Yet, direct measurements remain extremely scarce, covering less than 1% of marine fish species. Building on the Pace Of Life Syndrome (POLS) hypothesis, which posits a positive correlation between metabolism and life-history speed, we inferred temperature- and mass-specific resting metabolic rate for 18,214 fish species (16,998 Teleostei and 1,216 Elasmobranchii). Estimates were derived from relationships with 15 functional traits and phylogeny using phylogenetic structural equation modelling, a novel approach combining phylogenetic comparative methods with structural equation models. Cross-validations indicated high predictive performance, although some clades require cautious interpretation due to limited empirical data. In line with expectations from life-history theory, ecological niche and morphology, species with the highest metabolic rates were small, narrow, shallow-bodied, and long-jawed pelagic fish with high mortality, short lifespan and rapid growth — traits typical of small epipelagics. The metabolic rate variation aligned with the slow–fast life-history continuum, supporting the POLS hypothesis, but was equally explained by reproductive strategy: highly fecund, fast species exhibited the highest metabolic rate independent of mass and temperature, whereas low-fecundity slow species had the lowest. By filling a critical physiological data gap, our dataset provides a foundation to test long-standing ecological and evolutionary hypotheses such as POLS, and offers a powerful resource to improve models of species and community responses to climate change and exploitation, ultimately supporting physiology-informed ecosystem management.

## Introduction

Ocean warming and deoxygenation are intensifying and increasingly altering marine population dynamics (Calvin et al., 2023; Pinsky et al., 2020). Most research on climate change impacts on marine populations focused on latitudinal and depth shifts, altered phenology, and reduced body size (McKenzie et al., 2021; Pinsky et al., 2020). These observable consequences are largely explained by physiological processes (Jutfelt, 2020; Townhill et al., 2017).

Metabolic processes — the physiological and biochemical fluxes of material and energy within individuals (Brown et al., 2004) — play a central role in determining whole-organism responses to increased temperatures, salinity, or hypoxia (Nagelkerken et al., 2023). Resting metabolic rate (RMR) is the minimum metabolic activity required for an organism’s maintenance, typically measured in the laboratory on a resting organism (White and Marshall, 2023). Energy in excess of maintenance requirements is dedicated to other processes such as growth, maturation, and reproduction. Because metabolic rates govern energy allocation, they lie at the core of climate-induced changes in fish life-history traits such as body size, maturation age, or lifespan.

An organism’s fitness — its ability to contribute offspring to the next generation — depends on its life-history strategy, i.e. its life-history trait combination shaped by the energy allocation trade-offs between survival, growth, and reproduction in a constrained environment (Stott et al., 2024). The fast-slow continuum of life-histories distinguishes between species that grow rapidly, reproduce early, and have short lifespan versus those that grow slowly, mature late, and live longer (Gaillard et al., 2016). These differences shape population responses to climate change: for instance, fast life-history fish species are 25% more likely to experience population declines for a 1°C warming (Wang et al., 2020).

The Pace Of Life syndrome (POLS) hypothesis extends this continuum by suggesting that life-history, metabolic, and behavioral traits co-evolve under shared selective pressures and energetic trade-offs (Dammhahn et al., 2018). In this framework, the fast-slow life-history continuum correlates with metabolic rates and patterns of energy investment in growth, reproduction, and maintenance, which in turn determines how organisms respond to environmental stressors like climate change (Auer et al. 2018).

In ectotherms such as fish, empirical support for the POLS hypothesis has grown but remains incomplete (Wong et al., 2021). Species with a slower pace of life-- are expected to exhibit lower RMRs. In fish, empirical studies indeed found resting and/or routine metabolic rate to be negatively correlated with age at maturity (Gravel et al., 2024), as well as with vertical habitat and depth, as pelagics are more active than demersal and benthic species and deep-water species have slower life-histories (Ikeda, 2016; Killen et al. 2016). Furthermore, RMR correlates positively with the growth performance index, an integrated measure combining the von Bertalanffy infinite length and growth coefficient to provide a standardized measure of growth rate (Wong et al., 2021). Together, these relationships indicate that metabolism underpins both life-history evolution and ecological specialization.

Understanding the eco-evolutionary consequences of climate change on marine fish populations and communities therefore requires a physiological perspective (Jutfelt, 2020). Yet, direct metabolic data remain scarce, hindering our ability to predict how fish communities may respond to environmental changes. To overcome this limitation, we propose leveraging the relationship between metabolic rates and life-history traits to infer missing metabolic data. Rich trait databases, such as FishBase (Froese and Pauly, 2000), provide valuable resources for this approach. This database references trait values for approximately 35,600 fish species, with varying completeness among traits that clearly reflects the general lack of physiological data: while data on species habitat or trophic level are almost comprehensive, respiration data are available for only 423 species with respiration measurements from resting fish providing the closest proxy to RMR (Clarke and Johnston, 1999).

Here, we address the metabolic data gap using a novel approach, phylogenetic structural equation models (PSEM; Thorson et al., 2023) to infer missing RMRs. PSEM combine two complementary inference methods: phylogenetic comparative methods (PCM), which account for trait correlations among phylogenetically related species, and structural equation models (SEM), which specifies trait covariance structures among both quantitative and qualitative traits (Thorson et al., 2023; Thorson and van der Bijl, 2023). While phylogenetic imputation methods have already successfully generated large trait datasets for ectotherms — such as life-history traits for over 32,000 fish species using PSEM (Thorson et al., 2023) or metabolic index in marine ectotherms at smaller taxonomic scale using hierarchical phylogenetic factor analysis (Essington et al., 2024) — their potential remains underutilized for metabolic traits.

Our objectives are twofold. First, we use trait imputation based on PSEM to assemble a dataset of 16 traits for 18,214 fish species (16,998 Teleosteii and 1,216 Elasmobranchii), with a specific focus on inferring temperature- and mass-specific resting metabolic rate. Second, we use the inferred dataset to test predictions of the POLS hypothesis by examining the relationship between life-history strategies and resting metabolic rate. This approach aims to advance our understanding of how metabolism shapes life-history diversity and to provide a foundation for projecting the eco-evolutionary responses of marine fish communities to climate change.

## Material and Methods

### Metabolic and functional trait dataset

#### Resting metabolic rate data

RMR data were extracted from respiration measurements under controlled conditions, where RMR is traditionally estimated as the oxygen uptake of resting individuals (Clark et al., 2013; Chabot et al., 2016). Measurements were collected from five sources — the oxygen table of FishBase (Torres et Froese, 2000), Clarke and Johnston (2025), Gravel et al. (2024), Killen et al., (2016) and Ikeda (2016). RMR scales allometrically with body mass and follows the Arrhenius law in relation to temperature (Clarke and Johnston, 1999; Clarke & Fraser, 2004), expressed as:

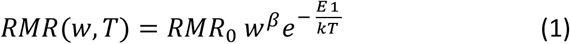

where *RMR*(w, T) is the resting metabolic rate (mgO_2_.h^-1^) of an individual of mass w (g) at temperature T (K), *RMR*_0_ the temperature- and mass-specific resting rate (mgO_2_.g^-^.h^-1^) the allometric scaling exponent with body mass, E the activation energy of the increase of RMR with temperature (eV), and k the Boltzmann constant in (eV.K^-1^). Details on the genus-level estimation of *RMR*_0_ are provided in the Supplementary Material (S1.a).

#### Functional trait data

16 traits across five categories (habitat, life history, morphology, diet, metabolism) were compiled from FishBase and complementary literature sources (Table 1; details in Supplementary Tables S1.b.1– S1.b.3) for 16,998 Teleosteii and 1,216 Elasmobranchii. Data cleaning, transformations, and filtering procedures are described in Supplementary Material (S1).

**Table 1.** Functional traits used in the PSEMs, with details on variable name, trait category (following Hadj-Hammou et al. (2021)), data type (continuous or categorical), transformation applied, levels for categorical traits, number of observed values across the 18,214 species in the dataset, and data sources.

| Name | Trait category | Continuous (C)<br>or<br>Categorical (F) | Transformation<br>(if continuous) | Levels<br>(if factor-valued) | Number of<br>observed<br>trait values | Source |
| --- | --- | --- | --- | --- | --- | --- |
| Habitat | Habitat: position in the water column following *FishBase* classification. | F |  | Pelagic,<br>Benthopelagic,<br>Demersal. | 18,214 | FishBase (Froese and Pauly, 2000) |
| Trophic level | Trophic ecology | C | Identity |  | 18,214 | FishBase (Palomares and Sa-a 2000, Sa-a et al. 2000) |
| Infinite length ($L_{\infty}$) | Growth: according to von Bertalanffy growth curve. | C | Natural log | | 1,169 | FishBase (Binohlan and Pauly 2000) |
| Infinite weight ($W_{\infty}$) | Growth: according to von Bertalanffy growth curve. | C | Natural log | | 1,613 | FishBase (Binohlan and Pauly 2000) |
| Length at maturity/Infinite length ($L_m/L_{\infty}$) | Life-history: Beverton-Holt ratio (Beverton et al., 1992). | C | Natural log | | 297 | FishBase (Binohlan and Pauly 2000) |
| Age at maturity, times mortality ($t_m * M$) | Life-history: Beverton-Holt ratio (Beverton et al., 1992).<br>Time-related | C | Natural log | | 926 | FishBase (Binohlan 2000) |
| Maximum age | Life-history. Time-related | C | Natural log |  | 913 | FishBase (Binohlan 2000) |
| Absolute fecundity | Reproduction: absolute fecundity. Number of eggs an animal produces during each reproductive cycle. Time related | C | Natural log |  | 1,138 | FishBase (Torres 2000) |
| Growth coefficient (K) | Growth: according to vonBertalanffy growth curve. Time-related | C | Natural log |  | 1,853 | FishBase (Binohlan and Pauly 2000) |
| Natural mortality rate (M) | Life-history. Time-related. | C | Natural log |  | 818 | FishBase (Binohlan and Pauly 2000) |
| Maximum body width | Morphometrics | C | Natural log |  | 3,502 | Price et al., 2022 |
| Maximum body depth | Morphometrics | C | Natural log |  | 3,502 | Price et al., 2022 |
| Lower jaw length | Morphometrics | C | Natural log |  | 3,501 | Price et al., 2022 |
| Minimum caudal peduncle depth | Morphometrics | C | Natural log |  | 3,496 | Price et al., 2022 |
| Optimal temperature ($T_{opt}$) | Habitat | C | Identity | | 12,068 | FishBase (Froese and Pauly, 2000) |
| $RMR_0$ | Metabolism | C | Natural log | | 546 | FishBase (Torres et Froese, 2000), Clarke and Johnston (2025), Gravel et al. (2024), Killen et al., (2016), Ikeda (2016)<br>see S1.a for details |

Because *RMR*_0_ was estimated at the genus-level for 43 genera, we extended these values to the species level by assigning each genus-level estimate to all species within the same genus sharing the same vertical habitat as the species used for the estimation, a factor known to affect metabolic rates (Ikeda, 2016). As a result, 546 species were assigned an *RMR*_0_ value in the dataset.

To facilitate analysis, inconsistent values were removed and maturation age and length were replaced by Beverton-Holt life-history ratios (S1.b; Beukhof et al., 2019). These invariants, specifically the ratio of maturation length to infinite length and the product of maturation age and mortality rate, are more stable across species and provide standardized indicators more meaningful for interspecific comparisons (Beverton et al., 1992). Trait imputation with PSEM (see below) used these invariants, but the subsequent analyses used maturation traits back-transformed. Continuous traits were log-transformed to improve the linearity, except temperature and trophic level, which were already linearly related to other traits. The log-transformation was maintained throughout trait inference and subsequent analyses.

### Parameter estimation

#### Principles of phylogenetic structural equation models

PSEMs integrate SEMs with PCMs to analyze the relationships among species traits while accounting for shared evolutionary history (Thorson et al., 2023; Thorson and van der Bijl, 2023). SEMs model causal relationships among observed and latent variables, estimating both direct and indirect effects between quantitative and qualitative. However, because species traits are not independent due to phylogeny, SEM assumptions are violated. PSEMs address this by incorporating a phylogenetic correlation matrix among taxa for each trait derived from a phylogenetic tree, ensuring dependencies arising from shared ancestry are captured. Conversely, PCMs typically represent trait relationships through covariance matrices whose dimension scales quadratically with the number of traits, becoming quickly intractable and difficult to interpret. PSEMs mitigate this problem by simplifying the covariance structure through SEMs designed to depict a subset of potential variable connections based on theoretical or empirical knowledge, thereby limiting the matrix dimension and yielding path coefficients interpretable as partial regression coefficients. PSEMs can therefore analyze multiple interdependent quantitative and qualitative traits while accounting for phylogenetic non-independence.

A key feature, central to this paper, is PSEMs’ ability to infer missing traits values by leveraging both trait relationships and phylogenetic correlations, thus combining the strengths of SEMs and PCMs in this respect. Applying PSEMs involves three main steps detailed in the following subsections: (i) defining potentially several competing SEM structures relating traits, (ii) selecting an appropriate phylogenetic tree, and (iii) selecting the final SEM structure before inferring missing trait values. Full methodological details are given in Thorson et al. (2023) and Thorson & van der Bijl (2023).

#### Structural equation models

The SEMs used in this study were designed according to theoretical and/or empirical expectations with the objectives to (i) infer missing values of *RMR*_0_ and (ii) investigate its relationship with life-history strategies to test the POLS hypothesis. Accordingly, SEMs specified direct or indirect paths between *RMR*_0_ and functional traits, each path representing a plausible causal link.

Five SEMs were constructed from functional traits (Table 1, Fig. S2.1), grouped into two main categories (Clark, 2004):

1. Evolutionary models where variation in *RMR*_0_ arises as an evolutionary consequence of temperature and ecological selection pressures, with *RMR*_0_ influenced by multiple upstream traits without affecting downstream traits.
2. Mechanistic models, grounded in bioenergetics, where *RMR*_0_ is determined by morphometry and vertical habitat due to physical constraints on physiology (Koslow, 1996; Ikeda, 2016) and, in turn, influences life-history traits and trophic level, as both emerge from bioenergetics and body size (Andersen, 2019; Romanuk et al., 2011).

Mechanistic models were further divided into four variants, differing in their treatment of infinite size and temperature. Models 1-2 followed Thorson et al. (2023), treating infinite length and temperature as primary variables (Fig. S2.1; Palomares et al., 2022; Pauly, 1980). Models 3-4 regarded infinite length as a bioenergetic emergent property (Vandermeer, 2006), hence influenced by *RMR*_0_, while temperature affected only downstream traits (Fig. S2.1; Brown and Sibly, 2012). Models 1 and 2 differed in whether length influenced *RMR*_0_ directly or indirectly through morphology (red arrow, Mechanistic 2, Fig. S2.1; Rubio-Gracia et al., 2020), while models 3 and 4 differed in whether length replaced or preceded mass downstream of *RMR*_0_ (red text and arrow, mechanistic models 3 and 4, Fig. S2.1).

Each SEM had two variants: one where morphometry influenced only *RMR*_0_ except for peduncle depth, which also influenced trophic level under the assumption that it determines swimming speed (Koslow, 1996) and thus predation capacity, and one where all morphometric traits affected both *RMR*_0_ and trophic level under the assumption that they are all involved in predation ability (blue arrows, all models, Fig. S2.1; Price et al., 2019; Langerhans et Reznick, 2009). Hence, in total, ten SEMs were tested. Further rationale for traits links is provided in the Supplementary Material (S2).

#### Phylogenetic tree

PSEMs require a phylogenetic tree to represent species evolutionary proximity, as phylogenetic distances are key to accounting for non-independence in traits due to shared ancestry, which violates the assumption of SEMs. By incorporating these distances, PSEMs constrain differences in trait values between closely related species ensuring phylogeny is properly modeled (Thorson, 2023).

Because a unified molecular phylogeny linking Elasmobranchii and Teleostei is currently unavailable, we followed previous studies (e.g. Thorson et al., 2023; Essington et al., 2024) in using a taxonomic tree as proxy. We based our taxonomic tree on FishBase (Van Der Laan et al., 2014; Nelson et al., 2016) and completed missing family names with the World Register of Marine Species (WoRMS Editorial Board, 2024). Each species was classified by genus, family, order, and class. The taxonomy was then converted into a tree using the r-package *ape* (Paradis et al., 2024), with phylogenetic distances of 1 between successive taxonomic levels (i.e. from species to genus, genus to family, etc.), assuming Brownian motion for trait evolution.

Although less precise than a molecular phylogeny, the taxonomic tree retains a meaningful hierarchical structure to approximate evolutionary relatedness and enables inclusion of many species (18,214) while ensuring methodological consistency. A potential limitation of this choice is that taxonomic hierarchy may not always accurately reflect evolutionary divergence, particularly at deeper levels. However, given the scope of the study and the absence of a comprehensive phylogenetic alternative, utilizing a taxonomic tree remains a pragmatic approach.

### Model selection, predictive performance and missing data inference

We compared the PSEMs derived from the ten different SEM structures using two criteria: (i) the marginal Akaike Information Criterion (marginal AIC) and (ii) predictive performance for *RMR*_0_ using ten-fold cross-validations. The best PSEM, as determined by these criteria, was then used to impute missing *RMR*_0_ values. The sensitivity of inferred trait values to observed *RMR*_0_ data was estimated by Jackknife cross-validation.

Predictive performance, i.e., the accuracy of trait inference, was assessed through ten-fold cross-validation where a portion of observed data was withheld, the PSEMs fitted to the remaining data, and the withheld values inferred and compared to their known values. In each of the 10 iterations, 10% of species (without replacement) were assigned ‘NA’ for all their traits, and imputation was performed on this modified dataset (Fushiki, 2011). For continuous traits, inference accuracy was quantified by the Percentage of Variance Explained (PVE) relative to a null model (S3-Eq 3; Thorson et al., 2023), ranging from 0 (equivalent to null model predictions) to 1 (perfectly predictions). For categorical traits, inference accuracy was evaluated using the Area Under the Receiver Operator Characteristics curve (AUC), which measures classification performance between 0 and 1 for each boolean dummy variable representing a categorical trait level, accounting for false negatives and false positives (Thorson et al., 2023; r-package *pROC* v1.18.5, Robin et al., 2011).

Quantitative outliers in the cross-validation were identified by regressing inferred against observed trait values and calculating Cook’s distance for each point. Cook’s distances exceeding 4/n, with n the species number, identified outliers following Bollen et Jackman (1990). The analysis was conducted when the regression slope was close to 1, indicating an accurate inference model, as it was considered unnecessary otherwise.

Finally, the sensitivity of the selected PSEM outputs to observed *RMR*_0_ data was quantified with a Jackknife (leave-one-out) resampling procedure (Manly, 1997). Given 43 genus-level *RMR*_0_ initial estimates, 43 resampled datasets were utilised for the Jackknife. For each re-sampled functional traits dataset, the species’ *RMR*_0_ values of this genus were removed. Each dataset served to refit the PSEM and standard errors of path coefficients and species-level trait inferences were computed from the corresponding 43 outputs (S3-Eq 4).

### Output analyses

#### Relationships between resting metabolic rate, life-history traits, and functional traits

The SEM structure guided trait imputation by prescribing which path coefficients were included. From the fitted PSEMs, we extracted the estimated path coefficients (), with γ(B|A) denoting the expected change in trait *B* given a one-unit change in trait *A*, which is the partial contribution of *A* to *B* while accounting for other covariates. Because several traits were log-transformed (Table 1), path coefficients could not always be interpreted as linear relationships between the original untransformed traits; full interpretation of these transformation cases is provided in Supplement Table S2.2 (Lefcheck, 2021 n.d.).

A key strength of (P)SEMs is their ability to investigate both direct and indirect relationships between variables by computing direct and indirect path coefficients. Raw path coefficients preserve the original measurement scales and are thus useful to compare different SEMs and interpret the covariance among traits in their original units. Standardized path coefficients, in contrast, assess the relative influence of predictors on a response trait within a given SEM. Therefore, we focused on direct and indirect standardized path coefficients linking *RMR*_0_ with other functional traits to explore the role of metabolism in the POLS.

#### Characterizing the relationship between life-history strategy and resting metabolic rate

Time-related life-history traits (e.g. growth coefficient, maturation age, fecundity per reproductive event, lifespan) rely on assimilated energy allocation between maintenance, growth and reproduction and are thus mechanistically linked to *RMR*_0_ (Brown et al., 2004, Gaillard et al., 2016). The POLS hypothesis also suggests covariation among life-history, behaviour and metabolism due to correlational selection partly grounded in bioenergetic trade-offs (Dammhahn et al., 2018). We therefore tested whether inter-specific variability in *RMR*_0_ maps onto gradients of time-related life-history traits across our 18,214 species.

To identify such gradients, we first applied archetypal analysis, an unsupervised learning method that identifies k extreme trait combinations (‘archetypes’) shaping a convex hull of the multivariate trait space (*archetypes* package, Eugster and Leisch, 2009). Species positions between archetypes (their coordinates) reflect proximity to these extremes, thereby providing interpretable trait-based gradients. The number of archetypes *k* was determined using the elbow method on the residual sum of squares (Fig. S4.1; RSS; i.e., identifying the number *k* beyond which adding archetypes only marginally reduces RSS). Trait values were standardized and the best solution was selected from 30 replicate runs.

Archetypal analysis was conducted specifically on time-related life-history traits (Table 1), i.e., maturation age (back-transformed from the Beverton-Holt ratio), maximum age, absolute fecundity (considered to be a reproduction rate), growth coefficient, and natural mortality, to capture the slow-fast life-history continuum (Gaillard et al., 2016). To illustrate each archetype with representative taxa, we used simplex plots in which each species’ coordinates represent the relative contribution of each archetype to its trait combination.

To investigate how *RMR*_0_ varied along time-related trait gradients and among archetypes, we performed a Principal Component Analysis (PCA) on this subset of time-related traits to summarize their variation along two major axes (r-package *FactorMineR* v.2.8, Le et al., 2008). We then overlaid *RMR*_0_ values on the PCA plane as a heatmap by gridding the two-dimensional space and assigning each cell the median *RMR*_0_ value of species it contained while projecting archetypes onto the same plane (r-package *ggplot2* v3.5.1, Wickham 2016). Finally, to reduce noise and highlight the *RMR*_0_ gradient more clearly, we regressed *RMR*_0_ against the first two principal components and mapped the model-predicted *RMR*_0_ values onto the PCA plane.

## Results

### Model selection supports mechanistic relationships between resting metabolic rate and functional traits

Differences in marginal AIC (ΔAIC) between the best-performing PSEM and alternative models ranged from 0.5 to 3,161.1 (Table S2.1, Fig. 1, Fig. S3.1). Across all model types (evolutionary or mechanistic 1-4), variants where trophic level was explained by all morphometric traits (variant b) consistently outperformed those relying on peduncle depth alone (variant a), with ΔAIC values from 14.2 to 82.9. Among these, mechanistic models 3 and 4 performed poorest (ΔAIC>690), reflecting excessive complexity relative to fit. The remaining models (evolutionary and mechanistic 1 and 2) all included temperature and body size as master variables, underscoring their central role in shaping functional strategies.

**Figure 1.**
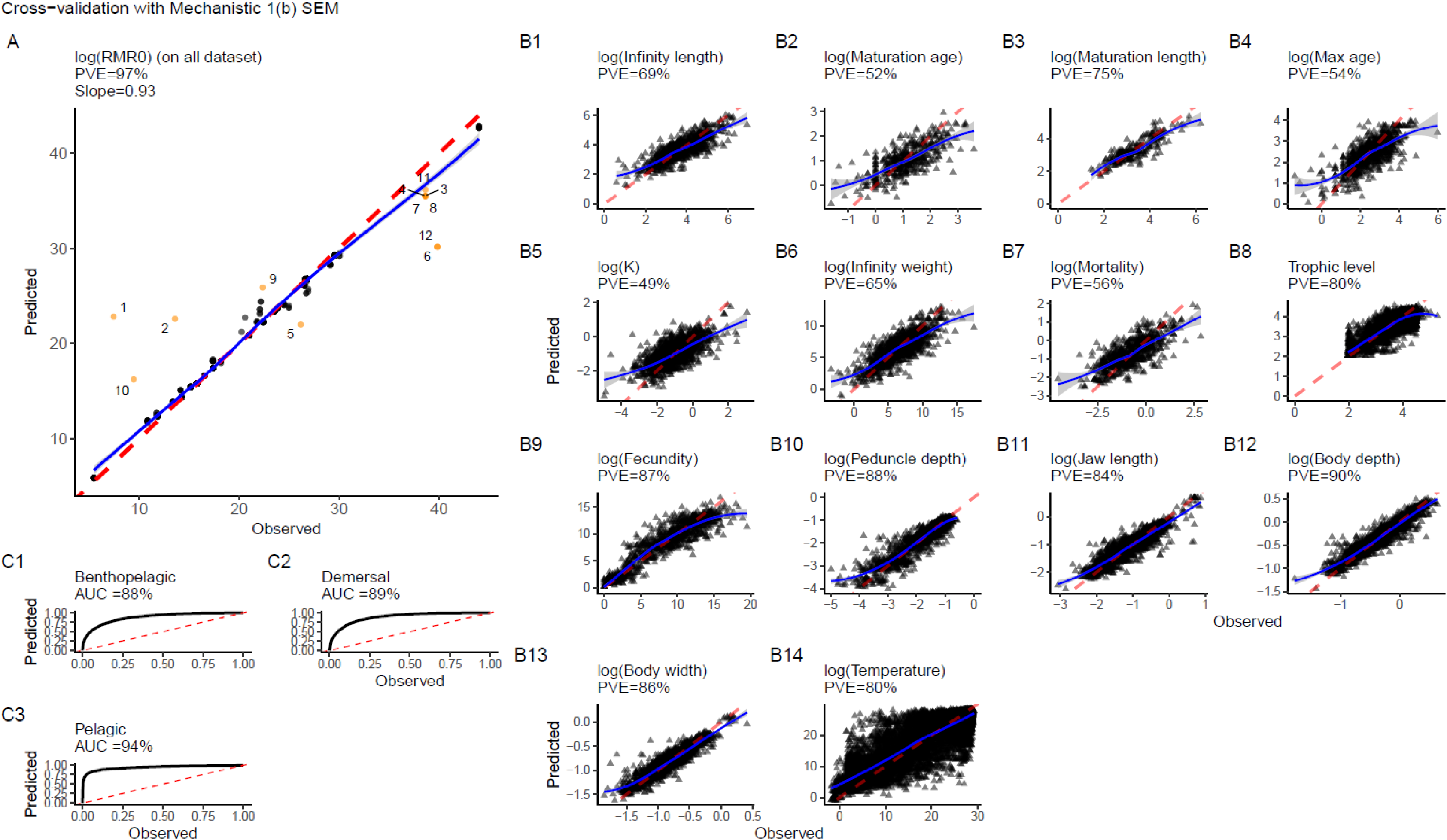
Ten-fold cross-validation results comparing inferred and observed trait values across the full dataset. Quantitative and qualitative cross-validation plots are shown separately. For quantitative traits (A and B), each point represents a species, the red dotted line is the 1:1 relationship, and the blue line shows a Loess regression. Inference quality increases as points approach the 1:1 line and PVE approaches 100%. A: *RMR*_0_ cross-validation plot. Orange points (numbered) are outliers. Corresponding species: (1) Channa brunnea, (2) Chaenocephalus aceratus, (3) Pseudopleuronectes obscurus, (4) Pseudopleuronectes yokohamae, (5) Chanos chanos, (6) Gillichthys seta, (7) Pseudopleuronectes americanus, (8) Pseudopleuronectes herzensteini, (9) Gilchristella aestuaria, (10) Mugil capurrii, (11) Pseudopleuronectes schrenki, (12) Gillichthys mirabilis. B: Other quantitative traits cross-validation plots. C: Qualitative traits cross-validations: vertical habitat binary variables. The black curve represents the Receiver Operator Characteristics curve (ROC). Classification quality increases as the ROC curve approaches the top-left corner and AUC approaches 100%.

Among these, bioenergetic-based models (mechanistic 1 and 2; ΔAIC of 0 and 0.5, respectively) outperformed the evolutionary model (ΔAIC = 16.1), which treated *RMR*_0_ variation as an outcome of natural selection by temperature and ecological processes through dependent functional traits. As ΔAIC ≤ 0.5 indicates equivalent support for competing models (Burnham and Anderson, 2002), we selected the simpler mechanistic model 1. This model included three master traits, temperature and body length, which influenced *RMR*_0_ indirectly through morphometry, and habitat, which affected *RMR*_0_ directly. *RMR*_0_, in turn, influenced bioenergetic fluxes and functional traits either directly (growth coefficient, mortality rate, infinite weight) or indirectly (trophic level, fecundity, maturation invariants, maximum age).

Ten-fold cross-validation of all variant b PSEMs confirmed that AIC-based model selection aligned with predictive performance. Overall, cross-validation results of the alternative models (Fig. S3.1) were similar to those of the best model (Fig. 1), confirming strong predictive performance of the method Quantitative traits other than *RMR*_0_ (Fig. 1B and Fig. S3.1B) had PVE values of 49 to 90%, with particularly high accuracy for morphometry, trophic level, fecundity and temperature (PVE ≥ 80%). Performance for habitat categories, the only qualitative trait, was also high, with AUC values of 88 to 94%, indicating high discriminative power (Fig. 1C and Fig. S3.1C). These results confirmed that the selected model reliably captured trait relationships as well as the quality and robustness of the inferred dataset for further analyses.

### Cross-validation reveals accuracy and limitations of resting metabolic rate imputation across fish species

The ten-folds cross-validation (Fig. 1A) showed low bias in predicted *RMR*_0_ values, with a slight tendency to underestimate high values and overestimate low values. The PVE was high (98%) and only 12 inferred outliers among 546 *RMR*_0_ values (Fig. S3.2), including eight overestimates and four underestimates.

Most outliers resulted from inference errors related to taxonomic biases within families or orders exhibiting high intra-group variability in *RMR*_0_, where closely related species had contrasting *RMR*_0_ values. For example, two outliers belonged to the Gobiidae family, which includes *Gillichthys* species with high observed *RMR*_0_ (log(*RMR*_0_)= 43.8 expected / ∼34.1 inferred) contrasting with *Glossogobius* species low observed values (log(*RMR*_0_) = 10.83 expected / ∼11.8 inferred). Consequently, *Gillichthys* species values were underestimated, whereas *Glossogobius* species values were overestimated, reflecting the PSEM’s tendency to reduce within-clade trait variability. A similar pattern occurred in Pleuronectiformes where *Pseudopleuronectes* species with high observed *RMR*_0_ (log(*RMR*_0_) = 38.62 expected / 35.50 inferred) were underestimated due to low observed values in *Pleuronectes* species (log(*RMR*_0_) = 22.19 observed / ∼22.4 inferred). Other cases of phylogenetic bias included Chaenocephalus aceratus (log(*RMR*_0_)≈ 13.59 expected / 22.59 inferred) and Gilchristella aestuaria (log(*RMR*_0_)≈ 22.37 expected / 25.88 inferred), both overestimated due to related species with higher *RMR*_0_. In contrast, inference error in *Chanos chanos* (log(*RMR*_0_)≈ 26.16 expected / 21.98 inferred), *Channa brunnea* (log(*RMR*_0_)≈ 7.44 expected / 22.83 inferred), and *Mugil capurii* (log(*RMR*_0_)≈ 9.46 expected / 16.24 inferred) could not be attributed to phylogenetic bias

The Jackknife cross-validation indicated low sensitivity of inferred trait values (excluding two traits without missing values: trophic level and habitat) to genus-level *RMR*_0_ data, with 91% of relative standard errors (RSEs) below 10% and 68% below 1%. Median RSEs ranged from very low to moderate: <1% for nine of the 14 inferred traits, 1-5% for four traits, and 11.5% for *RMR*_0_ itself (Fig. 2; Table S3.1). Despite the higher median RSE, *RMR*_0_ uncertainty remained moderate, with 75% of species below 16.7%, reflecting good precision despite limited empirical coverage in some lineages.

**Figure 2.**
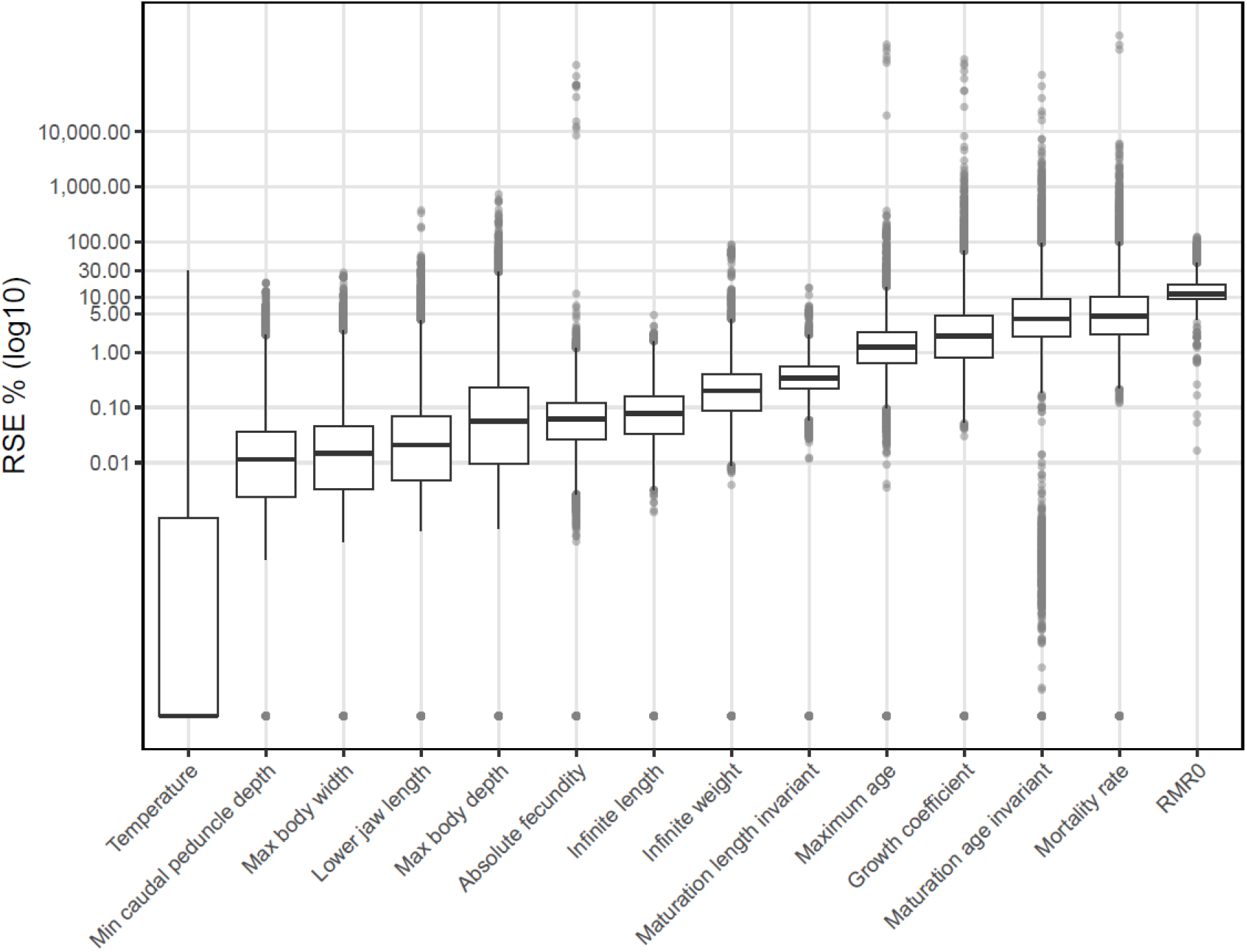
Variation in the Jackknife-derived relative standard errors (RSE) for each trait. The RSE values are log10-transformed. Boxplots show the median, interquartile range and twice the interquartile range (whiskers) of species RSE values associated with each trait. Each point represents the RSE of an outlier species.

Overall, ten-fold and Jackknife cross-validations indicated high predictive performance. These results confirm that the selected PSEM reliably captured metabolic patterns across taxa, although some *RMR*_0_ estimates should be interpreted cautiously within clades with limited observed data.

### The pace of life syndrome is well reflected in the structural equation model

The selected PSEM quantified which traits influenced or were influenced by others, including *RMR*_0_, through path coefficients (Fig. 3: direct paths only, Table S2.2: both direct and indirect paths). Traits affecting *RMR*_0_ varied in standardized path coefficient strength. Temperature ( ≈ −0.005) and infinite length ( ≈ 0.027) via indirect paths, and peduncle depth ( ≈ −0.049) via direct paths had negligible effects. Four traits showed stronger direct links to *RMR*_0_, including three morphometric traits: shallower body depth ( ≈ −0.181), narrower body width ( ≈ −0.085) and longer lower jaw length ( ≈ 0.097) were associated with higher *RMR*_0_. The relationships with vertical habitat matched life-style expectations: higher *RMR*_0_ was linked to pelagic habitat ( ≈ 0.141), typical of an active lifestyle, whereas demersal ( ≈ −0.118) and benthopelagic (γ≈ −0.051) habitats, characteristic of more sedentary lifestyles, were related to lower *RMR*_0_. Hence, pelagic, narrow-, shallow-bodied and long-jawed species exhibited higher *RMR*_0_, while temperature and body length had little influence.

**Figure 3.**
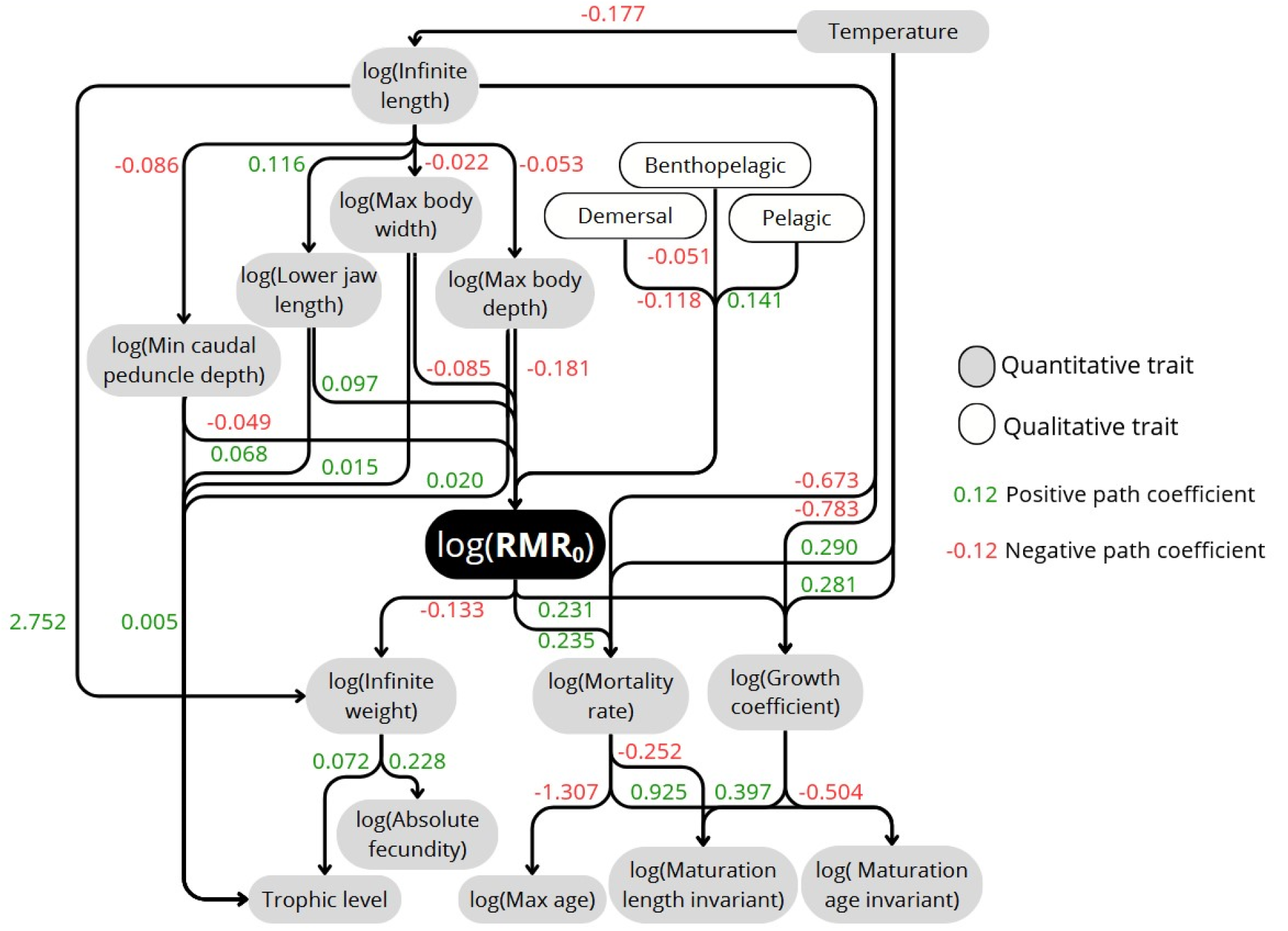
Selected structural equation model (SEM) to link the log of the temperature- and mass- specific resting metabolic rate (log(*RMR*_0_)) to the other functional traits. The color of the boxes differentiates quantitative traits (grey) and categorical traits (white). An arrow from box A to box B represents a causal relationship between traits A and B, each supported by a mechanistic link. The path coefficient associated with each arrow is the standardized partial correlation coefficient, colored red for negative and green for positive relationships. Indirect path coefficients between log(*RMR*_0_) and other traits are given in the supplementary material (Table S2.2).

Regarding *RMR*_0_ influence on other traits, two groups emerged. *RMR*_0_ had little effect on fecundity ( ≈ −0.030), trophic level ( ≈ −0.010) and the maturation length invariant ( ≈ 0.033) through indirect paths. In contrast, time-related life history traits were clearly influenced: higher *RMR*_0_ increased growth coefficient ( ≈ 0.231) and mortality rate ( ≈ 0.235) directly, and reduced maximum age ( ≈ −0.301) while increasing maturation age invariant ( ≈ 0.10) indirectly. Higher *RMR*_0_ was also directly associated with lower infinite weight ( ≈ −0.133). The unexpected positive correlation between *RMR*_0_ and the maturation age invariant resulted from the high variance in mortality rate, which masked the negative covariance between maturation age and *RMR*_0_. In summary, globally consistent with the POLS hypothesis, higher *RMR*_0_ correlated with a faster pace of life, characterized by higher mortality, shorter lifespan and faster growth towards asymptotic size, and a lower infinite weight, while trophic level and fecundity were only weakly affected.

### The temperature- and mass-specific resting metabolic of a species is linked to its pace of life syndrome but also its reproductive strategy

The archetypal analysis on time-related life-history traits identified three archetypes (Fig. S4.1) representing extreme pace-of-life strategies (Fig. S4.2): “fast” archetype, combining the highest growth coefficient and mortality rate with the youngest maximum and maturation ages and an intermediate fecundity; “slow” pace-of-life archetype, with the oldest maximum and maturation ages, and the lowest fecundity, growth coefficient and mortality rate; and “intermediate” archetype with intermediate values for most traits but the highest fecundity. These extreme archetypes thus aligned with the expected slow-fast life-history continuum, fast, slow and intermediate strategy, the latter distinguished by extreme fecundity. Representation in a ternary simplex plot showed that the small short-lived Engraulidae exemplified the fast archetype, long-lived Rajiformes the slow one, and Sparidae the intermediate one (Fig. 4).

**Figure 4.**
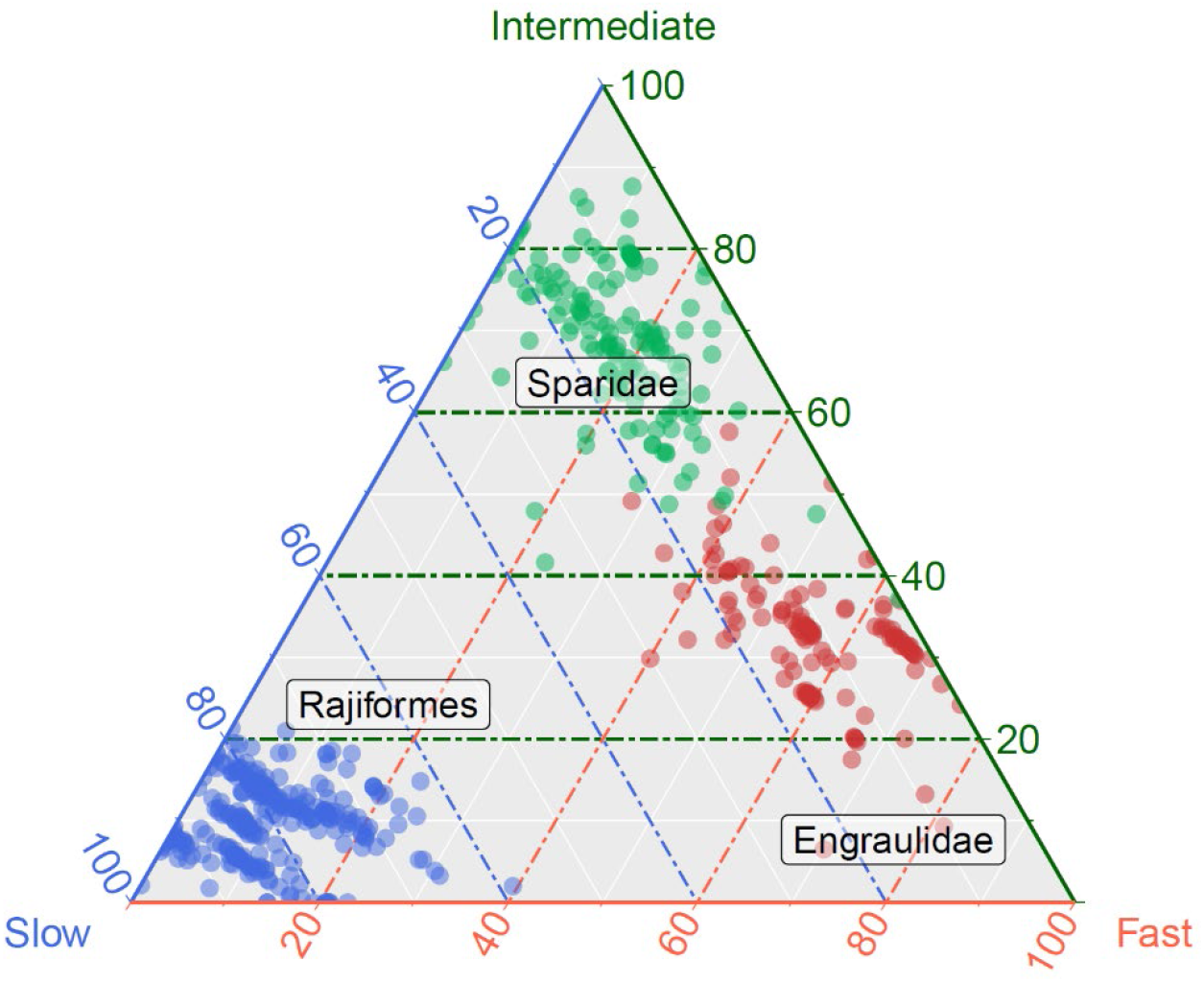
Ternary plots representing three functional trait groups identified through archetypal analysis (Eugster and Leisch, 2009). Each functional trait group is located at the apex of the triangles, and species (points) are classified along the gradients between archetypes. The archetypal groups range from fast to slow life cycle, encompassing fast, slow and intermediate species. One representative family or order is shown for each archetype. Point colors correspond to each archetype: fast (red), slow (blue), intermediate (green).

The two first principal components (PC1 and PC2) of time-related life-history traits explained 96% of their variance (Fig. 5.A). PC1 captured life-cycle speed variation (mortality rate, growth coefficient, maximum age, and partly maturation age) while PC2 reflected variation in reproductive strategy (fecundity and, partly again, maturation age). Hence, PC1 differentiated the fast (red triangle) from the intermediate (green circle) and slow (blue square) archetypes, while PC2 separated the slow from the intermediate archetype.

**Figure 5.**
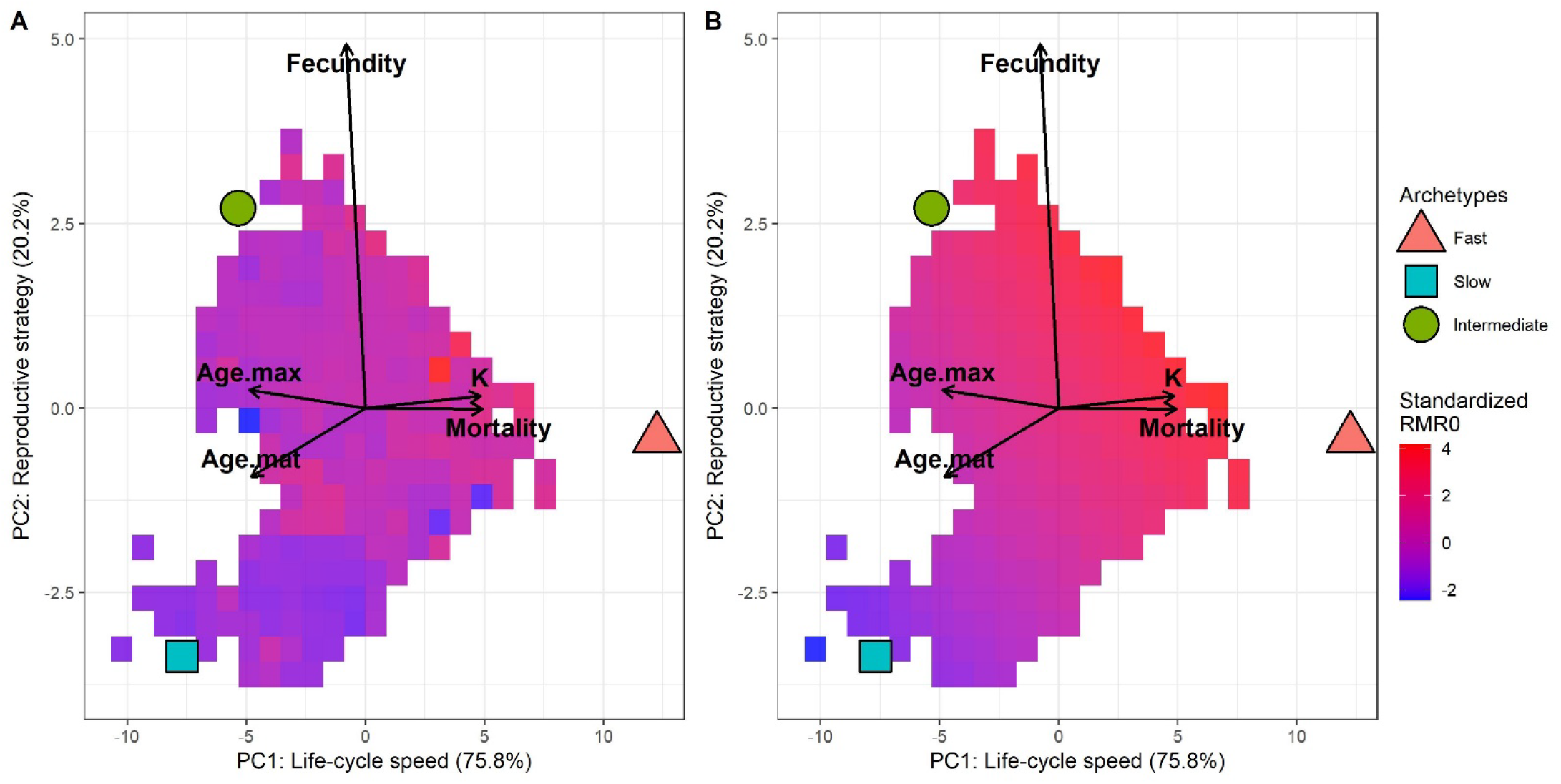
First and second principal components of the PCA based on life-history traits of fish with a complete inferred dataset. The traits included in the PCA are the maximum age (Age.max), the maturation age (Age.mat), the absolute fecundity (Fecundity), the growth coefficient (K) and the mortality rate (Mortality). Red triangles, blue squares and green circles represent fast, slow, and intermediate life-history strategies, respectively. Each square represents an aggregate of species with colors corresponding to the median species *RMR*_0_: A- estimated directly from the dataset, B- estimated from species’ *RMR*_0_ regressed and predicted against the two principal components.

Despite some variability, *RMR*_0_ gradients across life-history strategies followed POLS expectations, with median *RMR*_0_ increasing as life-history shifted towards a faster pace (Fig. 5.A). The steepest positive gradient ran from the slow to the fast archetype, with consistent shallower positive gradients from slow to intermediate and from intermediate to fast archetypes. Regressing and predicting *RMR*_0_ against the two PCs facilitated interpretation by removing variability unexplained by time-related life-history traits (Fig. 5.B, Table S4.1). The association of *RMR*_0_ with the reproductive strategy axis was stronger (PC2: 0.29, column Estimate Table S4.1) than with the life-cycle speed axis (PC1: 0.13) and the interaction term was negligible (0.01). However, when accounting for the variance explained by each axis, their relative importance was almost balanced: life-cycle speed contributed 46% and reproductive strategy 53%. This indicates that although the reproductive strategy axis is more strongly correlated with *RMR*_0_, the life-cycle speed axis remains nearly as influential in explaining predicted variability. Consequently, the strongest *RMR*_0_ gradient was diagonal starting from the slow archetype towards life-histories located between intermediate and fast archetypes.

## Discussion

In this study, we inferred a global metabolic dataset for marine fishes, providing temperature- and mass-specific resting metabolic rates *RMR*_0_, or maintenance costs in bioenergetic terms, for 18,214 species across 463 families, covering roughly 60% of marine fish diversity (Helfman et al. 2009). This addresses the general lack of physiological data in fish, considering that only ∼1.3% of species have metabolic rate records in the global database Fishbase (Froese and Pauly, 2000). The inferred dataset quantifies the fish maintenance costs. Unobserved *RMR*_0_ values were estimated together with 15 functional traits, expected to act as either determinants or consequences of metabolism, using PSEM. This approach exploits interspecific variability in both quantitative and qualitative traits, drawing on (i) life-history trade-offs and ecological constraints, and (ii) the shared evolutionary history of closely related species. The high quality of *RMR*_0_ inference confirms that physiological traits can be reliably estimated based on phylogeny and trait correlations.

### Reliability and limitations of trait inferences

The original dataset contained observed *RMR*_0_ for only 3% of species, with the remaining 97% inferred. This limited coverage could have biased estimates of *RMR*_0_ and correlated traits. Yet, high PVE and low jackknife RSE for strongly correlated traits — *RMR*_0_, body depth, growth coefficient, mortality, and infinite weight — in cross-validation indicate that the scarcity of *RMR*_0_ observations had minimal impact. Most importantly, the very high PVE and moderate jackknife RSE for *RMR*_0_ confirm that resting metabolic rate can be accurately estimated from correlations with functional traits and phylogenetic relationships.

The consistency between PSEM path coefficients and expected trait correlations across life-history, ecological, and morphological dimensions further supports the robustness of our inference. Species with the highest *RMR*_0_ were predicted to be small (low infinite weight), elongated (narrow-, shallow-bodied and long-jawed) pelagic fish characterized by high mortality, short lifespan and fast growth towards infinite length — traits typical of small epipelagics such as anchovies, sardines, herrings, mackerels and sprats (Ikeda, 2016; Wong et al., 2021). Path coefficients also aligned with the Beverton-Holt life history invariant hypothesis, which suggests that certain life-history trait ratios, such as mortality rate to growth coefficient, or products, such as mortality rate times maximum age, remain relatively constant across fish species (Beverton, 1992). Accordingly, *RMR*_0_ path coefficients with mortality, growth coefficient (positive) and maximum age (negative) had similar absolute magnitude. Furthermore, body size-trait relationships expected from bioenergetics and life-history theory were recovered, with larger body size associated with higher fecundity and lower mortality and *RMR*_0_ (Sentis et al., 2024). Together, these patterns also highlight the pivotal role of metabolism in balancing energetic investment among reproduction, growth, and survival.

Beyond life-history, metabolic differences also matched expectations across vertical habitats. Pelagic habitat was associated with higher *RMR*_0_, consistent with comparative studies showing elevated metabolic rates in pelagic versus demersal or benthopelagic fish, attributed to greater energetic costs of swimming and active lifestyle (Killen et al., 2016). Similar intraspecific patterns occur in *Perca fluviatilis*, where pelagic ecotypes have higher standard metabolic rates than littoral conspecifics (Andersson et al., 2022). Generally, metabolic rates decline with increasing depth across marine taxa, including fish, reflecting reduced locomotory activity and predator–prey interactions in dim-light environments (Seibel & Drazen, 2007). However, vertical habitat and depth are not equivalent: demersal species may inhabit shallow coastal areas, while pelagic species may occupy deep environments, explaining why bathypelagic fishes exhibit low *RMR*_0_ despite their pelagic lifestyle (Ikeda, 2016).

Similarly, morphological traits showed consistent relationships with metabolism, with higher *RMR*_0_, in species having narrow and shallow bodies, elongated jaws, and, to a lesser extent, narrow peduncles. Previous findings suggest such morphologies support faster swimming and active foraging, such as pursuit or cruise predation, elevating maintenance costs relative to deeper-bodied species (Videler, 1993; Killen et al., 2016). Jaw morphology further influences feeding mode and locomotion: elongated lower jaws enable wider gape and favor active predation such pursuit or ram feeding, which requires sustained swimming and foraging activity and thereby increased basal energetic costs compared to species with shorter jaws adapted to suction or ambush predation. These results highlight how morphology shapes baseline metabolic demand through its effects on locomotion, predation and baseline energy expenditure. Notably, morphological traits also influenced trophic level and enhanced model likelihood, leading to the selection of the Mechanistic model 1 (b). This underscores the importance of adequately representing non-focal traits in PSEMs, even when the focus is on inferring a specific trait such as *RMR*_0_.

Phylogenetically-informed imputation has previously provided robust inferences for large fish trait datasets (Thorson et al., 2023; Essington et al., 2024) and combining phylogeny with functional traits in statistical models generally improves explanatory power and inference performance (Thorson, 2023). However, evolutionary structure in PSEMs can also bias estimates. In our study, *RMR*_0_ outliers identified during cross-validation were often driven by closely related species, leading to over- or under-estimation inconsistent with their life-history strategies (Fig. S3.2). Based on their life-history traits combinations, most should have had average *RMR*_0_ values since they gathered near the center of the PCA biplot. Yet, 10 out of 12 observed *RMR*_0_ values fell outside the interquartile range (Table S1.b.2). These observations challenge the estimation process, because they were initially derived at the genus level from multiple species inhabiting the same habitat, and then assigned to species of the same genus and the same habitat in the dataset. Such group-level averaging inevitably obscures within-genus inter-specific variability. Restricting inference to the genus level would avoid this issue, but at the cost of sacrificing valuable species-level functional trait information. This highlights the need to expand species-level physiological observations to provide information on within-genus inter- specific variability. More generally, the PSEMs rely on maximum likelihood estimation, which tends to compress distribution tails and shrink parameters estimates towards the mean (James and Stein, 1961). These statistical properties should be considered for interpretation and future analyses.

### Variation in resting metabolic rate among marine fishes follows the pace of life syndrome hypothesis but also reflects reproductive strategy

Our results reveal that variation in *RMR*_0_ among marine fishes aligns closely with their position along the slow–fast life-history continuum (Figure 5B), supporting predictions of the POLS hypothesis (Dammhahn et al., 2018). Along this continuum, *RMR*_0_ followed a clear gradient from low to high values with increasing life-history speed. Long-lived species with slow growth to infinite size, delayed maturation, low fecundity, and low mortality — characteristic of the slow archetype — had lower average *RMR*_0_, whereas short-lived species with rapid growth, early maturation, average fecundity, and high mortality — typical of the fast archetype — displayed higher *RMR*_0_ on average. Species combining relatively slow growth and maturation with high fecundity — corresponding to the intermediate archetype — had intermediate average *RMR*_0_. Yet, median *RMR*_0_ varied around this gradient (Figure 5B) illustrating that metabolism constrains, but does not strictly dictate, life-history strategies, highlighting the interplay between energetic allocation rules and ecological contexts.

These metabolic–life-history associations correspond closely to the theoretical triad of strategies described by Winemiller and Rose (1992) and later refined by Pecuchet et al. (2017). The three archetypes — fast, intermediate, and slow — match the opportunistic, periodic, and equilibrium strategies, respectively: (i) opportunistic species invest energy in rapid growth and early maturation at a small size, at the expense of survival and fecundity; (ii) equilibrium species invest in survival, notably of juveniles through larger eggs and/or parental care, exhibiting slow growth, late maturation at large size, low mortality, and high longevity, at the expense of fecundity; and (iii) periodic species invest in fecundity, producing many small eggs and enhancing adult survival, at the expense of juvenile survival, with slower growth, later maturation at larger size, lower mortality and higher longevity than average (Figure 4 and S4.2). Our representative taxa fit these patterns: Engraulidae characterized by a fast life- cycle, Sparidae by high fecundity and a slower life-cycle, and Rajiformes by low fecundity and high juvenile survival, thanks to large eggs or ovoviviparity, and a very slow life-cycle.

However, the distribution of *RMR*_0_ across life-history strategies and the time-related life-history trait space indicates that basal metabolic diversity among fishes cannot be reduced to a single slow–fast continuum. A second dimension, related to reproductive strategy, explained *RMR*_0_ variation as much as life-cycle speed. Accordingly, the steepest gradient in *RMR*_0_ occurred between slow species and those positioned between the fast and intermediate archetypes, combining relatively rapid life cycles with higher fecundity than average (Figure 5B). This gradient, oriented at approximately 45° between the axes of life-cycle speed and reproductive strategy, highlights that both jointly shape metabolic demand. Such a pattern implies that maintenance costs integrate both temporal and reproductive components of energy allocation (Brown et al. 2004; Burton et al. 2011). From a bioenergetic perspective, supporting rapid somatic turnover and intense gamete production elevates baseline metabolism, whereas species investing in survival maintain lower maintenance demand. Hence, metabolism mediates a multidimensional trade-off among growth, reproduction, and survival rather than a single fast–slow axis (Killen et al., 2016; Dammhahn et al., 2018). This multidimensionality reconciles the POLS with the Winemiller–Rose framework, emphasizing that metabolic rate reflects not only the tempo of life but also the strategy by which energy is partitioned between fecundity and longevity. Understanding how these combined constraints shape species’ responses to environmental change will be key for predicting metabolic sensitivity and resilience in marine fish assemblages. Indeed, fast species such as small pelagics are known to respond rapidly to environmental variability, partly because of their higher sensitivity associated with tighter energetic margins is coupled with fast demographic turnover (Checkley et al. 2009; Planque et al. 2010; Sentis et al. 2024). The question remains how such energetic constrains will affect species with a slower turnover but higher fecundity under increasing environmental variability.

### Towards metabolism-informed ecosystem modelling

Alongside their ecological and evolutionary significance, our results have direct implications for ecosystem modeling, fisheries management, and climate change research. Incorporating metabolism into life-history allows more mechanistic predictions of species’ energy allocation under changing environmental conditions. Because *RMR*_0_ scales with mortality and growth rates as predicted by theory (Brown et al. 2004), integrating metabolic constraints into bioenergetic and size-based ecosystem models should improve forecasts of species’ productivity, resilience, and responses to exploitation. The strong alignment between fish resting metabolism and the slow–fast continuum provides a physiological basis to anticipate climate-driven shifts in community composition, as warming may favor fast-strategy species with higher maintenance costs (Daufresne et al., 2009; Ohlberger, 2013).

Ecosystemic models help predict how species and ecosystems respond to global changes such as climate change and overfishing. However, their parameterisation remains limited by data availability and uncertainty, which are often difficult to quantify (Heymans et al. 2020; Steenbeck et al. 2021). Currently, modelers collect parameters from literature or databases, and when species-specific values are missing, they are deduced from evolutionarily and/or ecologically close taxa relying on expert judgment. Although semi-automated methods have been proposed (e.g. Lovindeer et al. 2023), expert judgement alone remains insufficient for poorly documented parameters, such as metabolic traits. The *RMR*_0_ dataset inferred here offers a consistent and empirically grounded basis to constrain such parameters, linking species’ routine maintenance energy requirements to body mass and temperature. These data can directly inform bioenergetic frameworks, from single-species models (e.g. Dynamic Energy Budget; Kooijman, 2009), to multispecies ecosystem models (e.g. Bioen-OSMOSE, Atlantis; Audzijonyte et al., 2017; Morell et al., 2023) and global-scale models (e.g. APECOSM; Dalaut et al., 2025).

Together, these advances highlight the value of integrating physiological diversity and metabolic scaling into ecosystem assessments and conservation strategies, ensuring that metabolic constraints and adaptive potential are realistically represented in future projections of marine ecosystem dynamics.

### Physiological data needs and future directions to improve predictions

Further physiological data collection is essential to improve metabolic inference. At the order level, the heterogeneity of data availability is striking. Among elasmobranchs, metabolic data exist mainly for large sharks (51 species, 4 genus) but only for a single genus of rays (11 species). Among teleosts, 36% of Salmoniformes species have measurements, but none of the Carangiformes or Mulliformes species. Yet, Carangiform species such as *Xiphias gladius* and *Trachurus picturatus* are of major commercial importance; improved knowledge of their physiology and life history would therefore enhance model reliability benefiting fisheries management.

Metabolic parameters vary not only between but also within species. Thermal preferences differ across life stages: for instance, spawning adults and embryos have narrower thermal ranges than larvae and nonreproductive adults (Dahlke et al., 2020). Moreover, metabolic rates fluctuate through ontogeny with growth or food consumption (Rosenfeld et al., 2015), while mass-specific metabolic rates generally decrease with ageing (Glazier, 2022). Although our dataset can help parameterize species and ecosystem models, it represents adult routine maintenance rates. Extending coverage across life stages would directly improve the realism of population dynamic models and help identify energetically critical life stages, supporting fisheries management. Furthermore, metabolic rates depend on species and their functional traits but also their environment. For instance, a modelled cod population showed that under increasing fishing mortality, fitness optimization involved higher reproductive investment (Audzijonyte and Richards, 2018).

The interplay among metabolism, functional traits and environment is central to understanding fish resilience under global changes. Oxygen availability governs aerobic energy production, while warming simultaneously accelerates metabolism and exacerbates resource limitations (Nagelkerken et al., 2023). Ocean deoxygenation further constrains eco-evolutionary dynamics and ecosystems communities. Prior works link declining fish size with rising temperatures and decreasing oxygen concentrations (Audzijonyte et al., 2016). Our results add a physiological perspective: small opportunistic species incur the highest routine maintenance costs relative to mass and temperature. Their narrow energetic margins accelerate and amplify their responses to environmental variability, potentially destabilizing communities. Identifying such high-risk species will improve bioenergetic- based models’ projections to inform ecosystem and fishery management plans. Broadening and refining metabolic datasets across taxa and life stages will ultimately improve our understanding of fish resilience to global changes.

## Supporting information

Supp_Beneat_et_al_Filling_a_metabo

## Acknowledgements

This work was supported by the Horizon Europe research and innovation programme under grant agreement No 101060072 (ActNow), France Filière Pêche under grant agreement PH/2022/10 (ADAPT), and the Ifremer project FORESEA205. MB was supported by a PhD grant from Ifremer. The authors acknowledge the Pôle de Calcul et de Données Marines (PCDM, http://www.ifremer.fr/pcdm) for providing DATARMOR storage, data access, computational resources, visualisation and support services. YJS acknowledges support from the Pew fellows program in Marine Conservation at the Pew Charitable Trusts. We are grateful to J. Thorson for his guidance on the PSEM method and to L. Pécuchet for her advice on the archetypal analysis.

## Data availability statement

All data and code used in this study are publicly available. The inference process configuration for our analysis can be accessed at https://github.com/marinebnt/Mass_specific_maintenance_dataset_construction_and_analysis. The data generated during the data construction process, along with the scripts used to produce the figures in this paper, are also provided. All repositories are archived under the Zenodo record https://zenodo.org/records/17583188.

