## Supplementary material for "Filling a metabolism data gap in marine fish reveals dual pace-of-life and reproductive strategy axes Fish Metabolism and Pace-of-Life": Supp_Beneat_et_al_Filling_a_metabo

#### S1 - Supplementary material 1 - Data extracted as input for inference method

##### S1.a - Resting metabolic rate data

Experimental data on $RMR(w,T)$ were used to estimate temperature- and mass-specific resting metabolic rate $RMR_0$ and activation energy $E$, our metabolic traits of interest, by linearizing equation (1) as follows:

$$ln(\frac{RMR(w,T)}{w^{\beta}}) = ln(RMR_0) - \frac{E}{k}\frac{1}{T} \qquad (2)$$

and applying linear regression models.

$RMR(w,T)$ data were collected from five sources — FishBase (using the oxygen() function (Torres et Froese, 2000) of the R package *rfishbase 4.1.2* (Boettiger et al., 2012), interrogated on 25/09/2025), Clarke and Johnston (2025), Gravel et al. (2024), Killen et al., (2016) and Ikeda (2016) — and all units were standardized to $mgO_2.h^{-1}$. We removed duplicated data points when temperatures and weights were equal, or when the difference between oxygen consumption, weight and temperature was low to account for unit conversion bias. We included respiration data if they met four conditions: measurements had to (i) quantify standard/resting metabolism or routine metabolism metabolism (with routine and standard metabolism comprising respectively 3/5 and 2/5 of our dataset), but not active metabolism (Chabot et al., 2016, Killen et al., 2010), (ii) report the measured individual's body mass $w$ and water temperature $T$, (iii) be taken without applying stress, and (iv) be from fish species inhabiting brackish or marine waters.

Both measurements, resting and routine metabolic rates, are known to differ because the latter includes energy expenditure associated with spontaneous, small-scale movements that are excluded from the former (Chabot et al., 2016, Killen et al., 2010). We included the routine metabolic rate due to data scarcity, thus needed to debias $RMR_0$ estimate to represent temperature- and mass-specific resting metabolic rate. A measurement fixed effect was included in the linear regression to exclude the routine measurement effect from $RMR_0$ estimation.

Respiration data on multiple individuals (n>2) per species were scarce (n=121), preventing us from estimating species-specific $RMR_0$ and $E$ for a sufficient number of species. Instead, we decided to proceed with the estimation of $RMR_0$ and $E$ at the genus level, using a linear mixed effect model following equation (2). It included genus as a random effect acting on both the intercept, i.e. $ln(RMR_0)$, and the slope according to $-\frac{1}{kT}$, i.e. $E$, while assuming an allometric scaling exponent of $\beta = 3/4$ following Gillooly et al. (2001) and Clarke and Fraser (2004). Both the variance and co-variance between the intercept and slope random effects were included. The model was fitted using the R package lme4 (Bates et al., 2015) (Table 1). A linear mixed effect model across all genus was preferred over separate genus-specific standard linear models to better capture within-genus variation while constraining variability among genera to ensure realistic ranges of estimates (compared to Gillooly et al., 2001).

To obtain robust parameter estimation, the dataset was further filtered based on the following restrictions applied to each genus: (i) at least three measurements to ensure reliable estimation of genus-level random effects, (ii) a minimum of 5% variation in body mass among measurements to capture body mass scaling, (iii) a minimum temperature range of one degree Kelvin among measurements to quantify temperature dependence, and (iv) an increase of respiratory rate with body mass and temperature to ensure consistency with expectations from equations (1) and (2). The resulting filtered dataset included 2,501 resting metabolic rate measurements across four different classes (Elasmobranchii, Teleostei, Petromyzonti, Chondrostei), along with corresponding body mass and temperature data, covering 132 marine species across 72 genera. 2,361 measurements for 107 species and 86 genera were from Fishbase, 72 for 28 species and 22 genera from Clarke and Johnston (2025), 28 for 28 species and 17 genera from Gravel et al., (2024), 26 for 25 species and 17 genera from Killen et al., (2016), and 14 for 11 species and 9 genera from Ikeda (2016). The final genus-level estimates of $RMR_0$ and $E$ for 43 Teleostei and Elasmobranchii genera were obtained as the sum of the corresponding fixed effects and random effect deviations. Given the nearly-perfect correlation between $RMR_0$ and $E$ ($r = 0.99$), only $RMR_0$ was retained as a metabolic trait for subsequent analyses.

Table S.1.a.1 | Effect of temperature and genus groups on the oxygen consumption of fish. Parameter estimates of fixed and random effects from GLMM (Eq.2) are presented. Estimates of the random effects correspond to their estimated variance. Standard errors of the random effects were calculated as the square root of the diagonal elements of the variance-covariance matrix of the model (R package *merDeriv*, Wang 2018). Standard errors can be calculated because the confidence intervals of the random effects are approximately symmetrical. Genus-specific intercepts, slopes and their standard deviations are provided in Table S1.3.

| | Estimate | Std error | $\chi^2$ | Df | p-value |
|---|---|---|---|---|---|
| *Fixed effect* | | | | | |
| (Intercept) | 0.11 | 0.10 | | | |
| Temperature | 0.75 | 0.06 | 74.32 | 1 | <2.2E-16 |
| *Random effect (Genus)* | | | | | |
| (Intercept) | 0.58 | 0.10 | 168.43 | 2 | <2.2E-16 |
| Temperature | 0.14 | 0.06 | 860.98 | 2 | <2.2E-16 |

#### S1.b - Functional trait data

##### Background and summary

This dataset gathers functional traits of marine species based on data extracted from Fishbase and Price et al., 2022. Fishbase is a global information system on fish, including extensive data on fish biology, ecology, and their functional traits. The data extraction criteria are inspired by Beukhof et al., 2019. The overall goal of this data extraction was to collect traits relevant to fish metabolic activities in order to infer missing values using the phylogenetic structural equation modelling (see Main Text). This approach allows us to estimate species-specific maintenance needs and to relate functional trait groups to metabolic rates.

##### Spatial and temporal coverage

We applied no restriction on the spatial or temporal coverage of the data. Yet, freshwater species were excluded as we were only interested in marine species.

##### Taxonomic coverage

The dataset includes 18,214 species from 3,104 genus and 463 families across Elasmobranchii and Teleostei classes. 16,998 species are Teleostei and 1,216 are Elasmobranchii. Marine species are selected by excluding Freshwater species (‘Species’ table, ‘Fresh’, ‘Brack’ and ‘Saltwater’ columns). Migratory species predominantly marine are kept (‘Species’ table, ‘AnaCat’ column, amphidromous and anadromous, diadromous, oceanodromous categories).

##### TRAITS

We collected data on 16 functional traits, including 1 categorical trait (habitat), and 15 continuous traits. Trait values were extracted from FishBase using the R package rfishbase (version 4.1.2), except for morphometric traits that were obtained from the FishShapes database (v1; Price et al., 2022) and for the temperature- and mass-specific maintenance rate ($RMR_0$), which was estimated in this study (see Main Text). Missing values were voluntarily kept for subsequent imputation during the inference process. Trait values lower or equal to zero were treated as missing by transforming them into NAs as none of the traits included null or negative values in its natural range. All continuous traits were log-transformed to improve linearity of trait relationships, except for Temperature and Trophic level that were already linearly related to other traits. A summary of the dataset and its sources is provided in Table 2.

Table S1.b.1 | Number of available and missing values for each continuous and categorical trait at the species level.

| | **Available** | | | **Missing** | | |
|---|---|---|---|---|---|---|
| | All | Teleostei | Elasmobranchii | All | Teleostei | Elasmobranchii |
| Habitat | 18,214 | 16,998 | 1,216 | 0 | 0 | 0 |
| Trophic level | 18,214 | 16,998 | 1,216 | 0 | 0 | 0 |
| Infinite length | 1,169 | 1,045 | 124 | 17,045 | 15,953 | 1,092 |
| Infinite weight | 1,613 | 1,492 | 121 | 16,601 | 15,506 | 1,095 |
| Maturation length invariant | 297 | 249 | 48 | 17,917 | 16,749 | 1,168 |
| Maturation age invariant | 926 | 824 | 102 | 17,288 | 16,174 | 1,114 |
| Maximum age | 913 | 813 | 100 | 17,301 | 16,185 | 1,116 |
| Absolute fecundity | 1,138 | 859 | 279 | 17,076 | 16,139 | 1,064 |
| Growth coefficient | 1,853 | 1,701 | 152 | 16,361 | 15,297 | 1,064 |
| Mortality rate | 818 | 756 | 62 | 17,396 | 16,242 | 1,154 |
| Max body width | 3,502 | 3,502 | 0 | 14,712 | 13,496 | 1,216 |
| Max body depth | 3,502 | 3,502 | 0 | 14,712 | 13,496 | 1,216 |
| Lower jaw length | 3,501 | 3,501 | 0 | 14,713 | 13,497 | 1,216 |
| Min caudal peduncle depth | 3,496 | 3,496 | 0 | 14,718 | 13,502 | 1,216 |
| Temperature | 12,068 | 11,243 | 825 | 6,146 | 5755 | 391 |
| $RMR_0$ | 546 | 484 | 62 | 17,668 | 16,514 | 1,154 |

Table S1.b.2| Summary statistics and units of continuous traits in the initial dataset, following natural log transformation and prior to the inference process.

| | Min | 1st Quantile | Median | Mean | 3rd Quantile | Max | Unit |
|---|---|---|---|---|---|---|---|
| **Trophic level** | 1.98 | 3.24 | 3.43 | 3.45 | 3.70 | 5.29 | |
| **log(Infinite length)** | 0.59 | 3.15 | 3.65 | 3.71 | 4.27 | 6.91 | cm |
| **log(Infinite weight)** | -3.33 | 4.97 | 6.60 | 6.59 | 8.22 | 17.49 | g |
| **log(Maturation length invariant)** | 1.45 | 2.76 | 3.25 | 3.36 | 3.96 | 6.70 | cm |
| **log(Maturation age invariant)** | -2.50 | 0.65 | 1.10 | 1.06 | 1.61 | 3.59 | y |
| **log(Maximum age)** | -1.39 | 1.95 | 2.57 | 2.50 | 3.09 | 5.97 | y |
| **log(Absolute fecundity)** | 0 | 3.74 | 8.91 | 8.17 | 11.99 | 19.52 | Number of eggs |
| **log(Growth coefficient)** | -4.96 | -1.69 | -1.16 | -1.08 | -0.51 | 3.06 | $y^{-1}$ |
| **log(Mortality rate)** | -4.54 | -1.47 | -0.87 | -0.80 | -0.15 | 2.82 | $y^{-1}$ |
| **log(Max body width)** | -1.85 | -0.97 | -0.85 | -0.84 | -0.71 | 0.41 | mm |
| **log(Max body depth)** | -1.77 | -0.49 | -0.24 | -0.27 | -0.04 | 0.63 | mm |
| **log(Lower jaw length)** | -3.06 | -1.29 | -0.96 | -1.00 | -0.69 | 0.87 | mm |
| **log(Min caudal peduncle depth)** | -4.98 | -1.78 | -1.39 | -1.59 | -1.17 | -0.59 | mm |
| **Temperature** | -1.80 | 11.30 | 23.3 | 19.32 | 27.60 | 29.20 | °C |
| **log($RMR_0$)** | 5.47 | 15.77 | 17.41 | 20.62 | 26.81 | 43.99 | $mgO_2.g^{-\beta}.h^{-1}$ |

***Habitat***

The vertical habitat trait gives the species' position in the water column. It was retrieved from the ‘Species’ table (‘DemersPelag’ column) in rfishbase. To reduce the complexity of the inference process, we simplified habitat categories into 3 levels:

- Pelagic species, including: ‘pelagic’, ‘pelagic-oceanic’, ‘pelagic-neritic’, and ‘bathypelagic’;
- Benthopelagic species, including ‘benthopelagic’, ‘reef-associated’ (following Watanabe and Payne, 2023)
- Demersal species including ‘demersal’ and ‘bathydemersal’ species.

As a result, 2,654 species were classified as pelagic, 6,430 species as benthopelagic and 7,919 as demersal.

***Trophic level***

Species' trophic level corresponds to their position in the food web. It was extracted from the ‘Ecology’ table (‘DietTroph’ column, and ‘FoodTroph’ when data was missing) in rfishbase. The ‘Estimate’ table (‘Troph’ column) was also used to complete the dataset.

***Infinite length***

Infinite length is a result of the von Bertalanffy growth equation and represents the maximum asymptotic length a fish can measure. Values are estimated as the mean of the 'Loo' column of the 'Popgrowth' table, from the *rfishbase* package, and completed with the 'Linf' column of the 'Estimate' table. Infinite lengths were constrained to respect a ratio of LInfinite/LMaximum in between [0.66777, 1.3333] according to Fishbase recommendations ("Note that studies, where Loo is very different (+/- 1/3) from Lmax, are doubtful"). Linf smaller than Lmat were cleared.

***Infinite weight***

Infinite weight is a result of the von Bertalanffy growth equation and represents the maximum asymptotic weight a fish can reach. Values are estimated as the mean of the 'Woo' column of the 'Popgrowth' from the *rfishbase* package.

***Length at maturity***

The length at which the probability of being mature is equal to fifty percent. It is the mean of the values extracted for each species in the 'Popgrowth' table of the 'Lm' column with the *rfishbase* table. Later this data is converted as the ratio between Maturation length and Infinite length (Andersen, 2019). This ratio forces the inference of missing morphological data to be into more realistic ranges (LInfinite>Lmaturity).

***Age at maturity***

The age at which the probability of being mature is equal to 50%. It is the mean of the values extracted for each species in the 'Maturity' table, 'tm' variable. It is completed by the 'AgeMatMin' and 'AgeMatMin2' variables if needed. Later, it is converted into a ratio of the age at maturity times mortality (Andersen, 2019). This allows us to infer ages at maturation lower than maximum ages during the inference process.

***Max age***

Corresponds to the maximum age a fish can reach. It is extracted using the mean values from the *rfishbase* package, column 'tmax' of the 'Maturity' table. Maximum ages smaller than Maturation ages were cleared.

***Fecundity***

Absolute fecundity is the number of eggs an individual lays per spawning event. These trait values are coming from *rfishbase* table 'Fecundity', 'FecundityMean', or 'FecundityMin' and 'FecundityMax' if missing data.

***K***

K is a parameter in the von Bertalanffy equation, showing how fast an individual reaches its asymptotic size. In the dataset, it corresponds to the mean per species of the 'K' parameter from the 'Popgrowth' table of the *rfishbase* package.

***Mortality***

Mortality is the rate of deaths from various causes. In the dataset, it corresponds to the mean per species of the 'M' parameter from the 'Popgrowth' table of the *rfishbase* package.

***Body width: Max body width***

A fish's body width is the size of an individual when viewed from the front. Its values are extracted from the FishShapes v1 (Price et al., 2022) dataset. The value from the dataset was divided by the geometric mean, corresponding to the cubic root of the standard_length times max_body_depth times max_body_width from the same dataset.

***Body depth: Max body depth***

A fish's body depth is the height of a fish's body. Its values are extracted from the FishShapes v1 (Price et al., 2022) dataset. The value from the dataset was divided by the geometric mean, corresponding to the cubic root of the standard_length times max_body_depth times max_body_width from the same dataset.

***Jaw length: Lower jaw length***

Its values are extracted from the FishShapes v1 (Price et al., 2022) dataset. The value from the dataset was divided by the geometric mean, corresponding to the cubic root of the standard_length times max_body_depth times max_body_width from the same dataset.

***Peduncle depth: Min peduncle depth***

Its values are extracted from the FishShapes v1 (Price et al., 2022) dataset. The value from the dataset was divided by the geometric mean, corresponding to the cubic root of the standard_length times max_body_depth times max_body_width from the same dataset.

***Temperature***

Corresponds to the optimal temperature for an individual to live in. Extracted from FishBase as the 'TempPrefMean' of the rfishbase package, 'Estimate' function. If the value was 0, it was replaced by 0.1 for logarithmic conversion.

$$RMR_0$$

Mass-specific maintenance is the normalisation coefficient species-specific of the metabolic theory of ecology explaining maintenance rate. The details of this variable definition for the dataset used or the inference process are explained in the main text.

Table S1.b.3 | Genus-specific slopes and intercepts values with their standard errors from a linear mixed-effects model. Intercepts and slope were rescaled because the dataset was normalized for the analysis.

| | (Intercept) | Slope | Standard error Intercept | Standard error Slope |
|---|---|---|---|---|
| Abramis | 43.662064 | 12664.01165 | 0.32385424 | 0.10446117 |
| Acipenser | 20.661915 | 6258.36941 | 0.15562595 | 0.09422447 |
| Alburnus | 21.301889 | 6525.52662 | 0.30558656 | 0.32476643 |
| Ambassis | 11.077542 | 3862.09006 | 0.272991 | 0.27389063 |
| Ameiurus | 24.118786 | 7544.56348 | 0.12014282 | 0.12671603 |
| Anoplogaster | 26.708849 | 8239.43424 | 0.32553326 | 0.58745427 |
| Aphaniops | 15.979675 | 5020.68851 | 0.29239917 | 0.23156261 |
| Arnoglossus | 14.253345 | 4650.36449 | 0.30400232 | 0.30567155 |
| Brevoortia | 17.375815 | 5282.46279 | 0.15617879 | 0.13175453 |
| Buglossidium | 20.60062 | 6563.32626 | 0.30400232 | 0.30567155 |
| Callionymus | 26.808184 | 8112.24999 | 0.27559749 | 0.25495188 |
| Carassius | 19.002423 | 6022.11144 | 0.06431179 | 0.05807356 |
| Carcharhinus | 21.007855 | 6362.39788 | 0.23871634 | 0.28322822 |
| Catostomus | 14.199865 | 4578.30069 | 0.17190517 | 0.16270644 |
| Chaenocephalus | 13.594249 | 4221.81084 | 0.21726315 | 0.41056999 |
| Channa | 7.435901 | 2798.45731 | 0.12860614 | 0.16501568 |
| Chanos | 26.155167 | 7825.20963 | 0.2454287 | 0.28233107 |
| Chelon | 11.765208 | 3758.66 | 0.08167596 | 0.07636934 |
| Chiloscyllium | 18.158112 | 5748.08468 | 0.30783908 | 0.27644405 |
| Chromis | 15.766386 | 5011.40617 | 0.27383339 | 0.26107141 |
| Cirrhinus | 14.480567 | 4720.07845 | 0.10534168 | 0.12307967 |
| Colossoma | 22.904584 | 7033.09418 | 0.15295891 | 0.18651474 |
| Coregonus | -0.95869 | -93.82626 | 0.28362069 | 0.35845172 |
| Cyprinus | 10.629059 | 3453.20539 | 0.07037342 | 0.06015851 |
| Dorosoma | 21.798469 | 6396.79179 | 0.13349184 | 0.09117011 |
| Eleginus | 18.197838 | 5499.26994 | 0.2907479 | 0.45202337 |
| Engraulis | 43.993572 | 12501.34834 | 0.35773207 | 0.17606397 |
| Esox | 12.950119 | 4081.40224 | 0.14177242 | 0.15097483 |
| Exodon | 8.982661 | 3304.06925 | 0.2705269 | 0.33239081 |
| Gadus | 14.140829 | 4331.21198 | 0.0929289 | 0.12139432 |
| Gambusia | 20.101616 | 6318.97471 | 0.13133685 | 0.1432158 |
| Gasterosteus | 24.484857 | 7475.57811 | 0.30437866 | 0.34613693 |
| Gilchristella | 22.365075 | 6943.16667 | 0.21061277 | 0.12637663 |
| Gillichthys | 39.829038 | 12177.7138 | 0.14861956 | 0.06659806 |
| Glossogobius | 10.834003 | 3585.94176 | 0.25643072 | 0.23069811 |
| Gobio | 21.947637 | 6644.93264 | 0.32752765 | 0.23846984 |
| Heteropneustes | 13.084584 | 4353.66009 | 0.21566542 | 0.23432075 |
| Hippocampus | 15.184428 | 4904.72506 | 0.305476 | 0.24857217 |
| Hippoglossoides | 22.088359 | 6828.25762 | 0.30568876 | 0.61582294 |
| Labeobarbus | 21.212796 | 6628.77138 | 0.12316096 | 0.09644324 |
| Lampetra | 39.024767 | 11765.40423 | 0.16059453 | 0.17721086 |

| | | | | |
|---|---|---|---|---|
| Lepomis | 1.62852 | 819.58458 | 0.06701984 | 0.06392503 |
| Leuciscus | 30.632684 | 9129.33268 | 0.22518641 | 0.21137721 |
| Limanda | 26.767144 | 8030.63432 | 0.33429282 | 0.46846408 |
| Lipophrys | 26.925059 | 8130.96284 | 0.33060993 | 0.11420859 |
| Lutjanus | 17.405839 | 5414.28884 | 0.26604185 | 0.30205286 |
| Macrognathus | 10.813069 | 3729.18375 | 0.14045348 | 0.15681619 |
| Mugil | 9.457532 | 3066.918 | 0.09259062 | 0.07925484 |
| Mystus | 13.798135 | 4428.30246 | 0.12787445 | 0.18390751 |
| Notothenia | 13.387569 | 4174.01438 | 0.27039366 | 0.62745294 |
| Oncorhynchus | 26.548981 | 7953.53776 | 0.06712787 | 0.07209837 |
| Ophiodon | 20.250942 | 6165.03348 | 0.35907856 | 0.32775435 |
| Oreochromis | 25.50617 | 7922.72588 | 0.0684534 | 0.073488 |
| Perca | 22.538785 | 6895.3484 | 0.25987142 | 0.13866806 |
| Petromyzon | 21.765635 | 6676.62464 | 0.21437732 | 0.22058507 |
| Platichthys | 25.176966 | 7649.81616 | 0.13790852 | 0.13540597 |
| Pleuronectes | 22.18558 | 6771.82563 | 0.11137855 | 0.11680086 |
| Pomadasys | 29.542262 | 8858.58802 | 0.17183121 | 0.10762875 |
| Pseudopleuronectes | 38.623601 | 11630.48977 | 0.10506134 | 0.07316975 |
| Salmo | 26.311789 | 7948.33322 | 0.12800841 | 0.12933604 |
| Sander | 10.468201 | 3452.40738 | 0.19623959 | 0.14052609 |
| Scomber | 30.050111 | 8941.3224 | 0.25039659 | 0.24482041 |
| Scophthalmus | 23.871505 | 7247.06036 | 0.33667321 | 0.25703836 |
| Scyliorhinus | 5.466438 | 2056.21745 | 0.15911451 | 0.1219719 |
| Sebastolobus | 24.979754 | 7806.24981 | 0.33205255 | 0.55935215 |
| Solea | 23.546135 | 7148.89045 | 0.33572156 | 0.21719987 |
| Squalius | 24.046305 | 7286.79966 | 0.34873432 | 0.27828733 |
| Squalus | 22.413558 | 6745.97925 | 0.29606925 | 0.26829971 |
| Thunnus | 26.594542 | 7703.13688 | 0.26119648 | 0.23355214 |
| Torpedo | 16.549422 | 5246.65706 | 0.36016834 | 0.26819223 |
| Trematomus | 11.859537 | 3802.59348 | 0.26430411 | 0.6716183 |
| Zoarces | 29.113179 | 8801.40827 | 0.31970476 | 0.57488577 |

### S2 - Supplementary material 2 - Structural equation model (SEM)

S2| Supplementary material and methods: SEM assemblage explanation

Five structural equation models were assembled from the set of functional traits considered. The first model was based on evolutionary drivers, while the other four models were motivated by mechanistic drivers (see Main Text). The models shared two common features: they all explained $RMR_0$in relation to habitat and they all linked body size to fecundity, trophic level, K and mortality rate, as well as maturation invariants and maximum length. Additional evolutionary and mechanistic considerations were used to further explain the relationships among traits.

Habitat explains $RMR_0$ because deeper-dwelling fish have higher water content and lower maintenance metabolic rate (Torres et al., 1979; Ikeda et al., 2016).

The relationship between traits and body size was interpreted through bioenergetics principles. Absolute fecundity depends on asymptotic size, reflecting the allometric relationship between body mass and reproductive investment. Trophic level is strongly related to body size due to opportunistic predation.

There is an evolutionary trade-off between mortality and growth rate. The rate at which an individual approaches its asymptotic length (K) or dies influences the age at maturation, and consequently the maturation length and maximum age. Higher mortality rates are associated with shorter lifespans and earlier maturation.

Table S2.1 | AIC and ΔAIC values of the SEM models. Each row of the table represents a different SEM, (a) corresponding to the SEM where only the peduncle depth explains the trophic level, (b) corresponding to the SEM where only the morphometric traits explain the trophic level. The AIC in bold is the lowest, and this model was selected for further analyses. ΔAIC represents the difference in marginal AIC values (ΔAIC) between the best-performing SEM and alternative models.

| | AIC | ΔAIC |
|---|---|---|
| Evolutionary (a) | 113,494.9 | 30.3 |
| Evolutionary (b) | 113,472.7 | 16.1 |
| Mechanistic 1 (a) | 113,476.6 | 20.0 |
| Mechanistic 1 (b) | **113,456.6** | 0.0 |
| Mechanistic 2 (a) | 113,479.0 | 22.5 |
| Mechanistic 2 (b) | 113,457.1 | 0.5 |
| Mechanistic 3 (a) | 116,817.7 | 3,361.1 |
| Mechanistic 3 (b) | 116,784.1 | 3,327.6 |
| Mechanistic 4 (a) | 114,179.5 | 772.9 |
| Mechanistic 4 (b) | 114,146.5 | 690.0 |

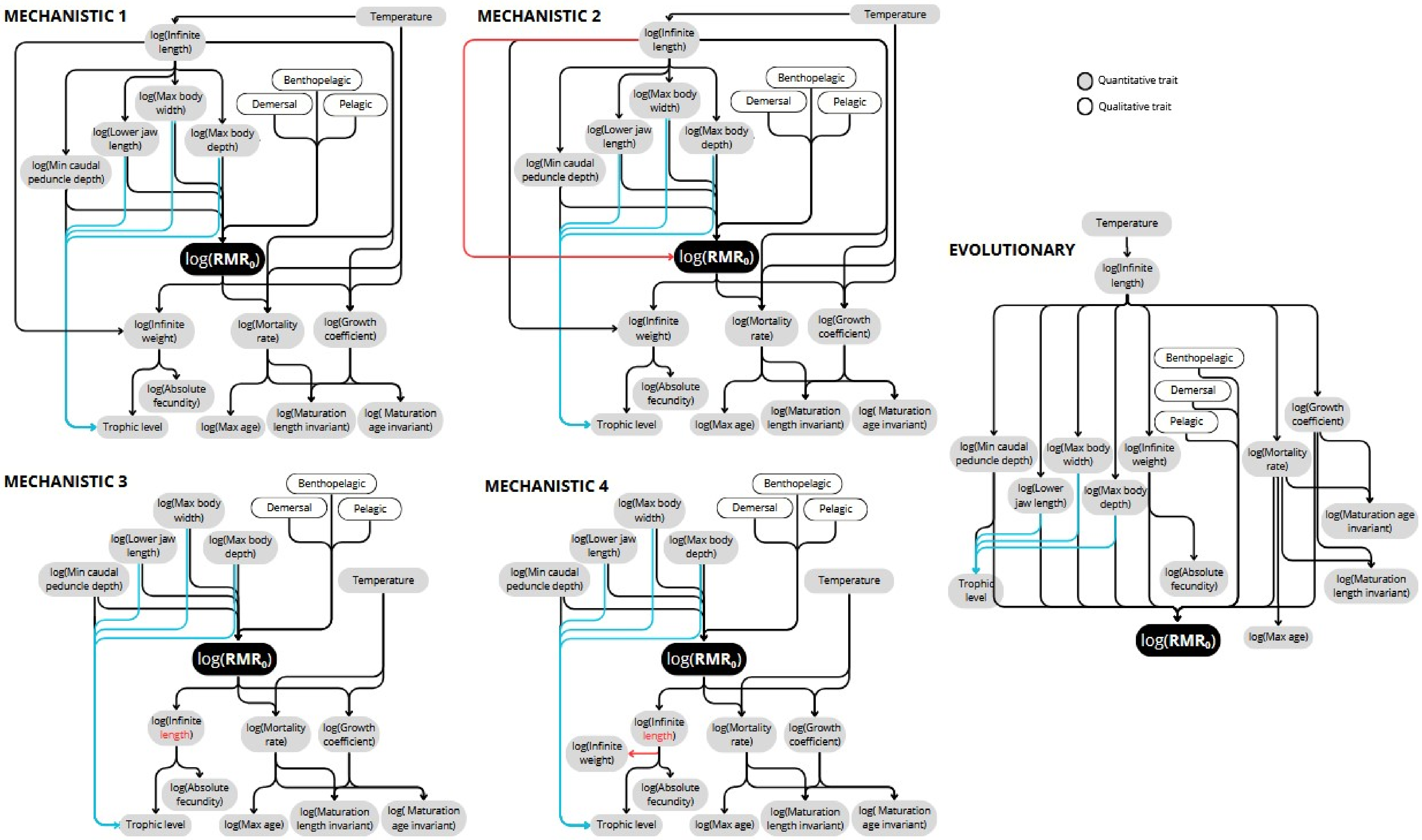
MECHANISTIC 1
Temperature
log(Infinite length)
log(Max body width)
log(Lower jaw length)
log(Max body depth)
log(Min caudal peduncle depth)
Benthopelagic
Demersal
Pelagic
log(RMR0)
log(Infinite weight)
log(Mortality rate)
log(Growth coefficient)
log(Absolute fecundity)
Trophic level
log(Max age)
log(Maturation length invariant)
log( Maturation age invariant)
MECHANISTIC 2
Temperature
log(Infinite length)
log(Max body width)
log(Lower jaw length)
log(Max body depth)
log(Min caudal peduncle depth)
Benthopelagic
Demersal
Pelagic
log(RMR0)
log(Infinite weight)
log(Mortality rate)
log(Growth coefficient)
log(Absolute fecundity)
Trophic level
log(Max age)
log(Maturation length invariant)
log( Maturation age invariant)
Quantitative trait
Qualitative trait
EVOLUTIONARY
Temperature
log(Infinite length)
Benthopelagic
Demersal
Pelagic
log(Growth coefficient)
log(Min caudal peduncle depth)
log(Max body width)
log(Infinite weight)
log(Mortality rate)
log(Lower jaw length)
log(Max body depth)
log(Maturation age invariant)
Trophic level
log(Absolute fecundity)
log(Maturation length invariant)
log(RMR0)
log(Max age)
MECHANISTIC 3
Benthopelagic
Demersal
Pelagic
log(Max body width)
log(Lower jaw length)
log(Max body depth)
log(Min caudal peduncle depth)
Temperature
log(RMR0)
log(Infinite length)
log(Mortality rate)
log(Growth coefficient)
log(Absolute fecundity)
Trophic level
log(Max age)
log(Maturation length invariant)
log( Maturation age invariant)
MECHANISTIC 4
Benthopelagic
Demersal
Pelagic
log(Max body width)
log(Lower jaw length)
log(Max body depth)
log(Min caudal peduncle depth)
Temperature
log(RMR0)
log(Infinite length)
log(Infinite weight)
log(Mortality rate)
log(Growth coefficient)
log(Absolute fecundity)
Trophic level
log(Max age)
log(Maturation length invariant)
log( Maturation age invariant)

Figure S2.1| Structural equation models (SEMs) structure. These SEMs were tested during the process of choosing a structural equation model connecting traits, to infer missing values of the $RMR_0$ parameter. Two categories of models were tested: (i) Evolutionary models, based on Thorson et al., 2023, and (ii) Mechanistic models, based on bioenergetic theories. Each model is read from top to bottom, following the arrows, when an arrow indicates the influence of one trait on another. Quantitative traits are shown in grey and qualitative (binomial) traits in white. Red arrows or red text highlight differences between the first mechanistic model (Mechanistic 1), and the other options, while blue arrows indicate alternative models. Each plot represents two SEM variants differing in how swimming speed is defined: variant a, based on peduncle depth only, or variant b, based on multiple morphometric traits.

Table S2.2 | Structural equation model path coefficients and their standard errors. Three categories of parameters are presented: (i) PC - direct path coefficients and their identification numbers, (ii) iPC - total path coefficients and their identification number (including only the total path coefficients towards or coming from $RMR_0$), (iii) V[trait] - trait variances. Two types of path coefficients are provided: raw (non-standardized) and standardized, with the latter recommended for comparing the relative effect of different variables. 'H:' denotes Habitat.

*__Notes on trait transformations__*: Path coefficients interpretation is affected by transformations, such as log-transformation applied to continuous traits (Table 1) (Lefcheck, 2021 n.d.). If the 'explanatory' trait is log-transformed as $A = ln(a)$, then a $x\%$ change in $a$ leads to a $ln(\frac{100+x}{100})\gamma(B|A)$ unit change in B, which approximates to a $x\gamma(B|A)\%$ change for small $x$ (as in such case $ln(\frac{100+x}{100}) \simeq \frac{x}{100}$. In contrast, if the 'explained' trait is transformed as $B = ln(b)$, then a $x$-unit change in $A$ leads to a $100(exp(\gamma(B|A)x) - 1)\%$ change in $b$. Lastly, if both traits are log-transformed, then a $x\%$ change in $a$ leads to a $100(exp(\gamma(B|A)ln(\frac{100+x}{100}) - 1)\%$ change in $b$, which approximates to $100(exp(\gamma(B|A)x) - 1)\%$ for small $x$.

| *Path* | *Parameter* | *Path coefficient* | *Standard error* | *Relative standard error (%)* | ***Standardized path coefficient*** |
|---|---|---|---|---|---|
| H:Benthopelagic -> log($RMR_0$) | PC0 | -0.068624445 | 0.086074981 | 125.4290382 | -0.051121357 |
| H:Demersal -> log($RMR_0$) | PC1 | -0.14992821 | 0.080875746 | 53.94298133 | -0.118364691 |
| H:Pelagic -> log($RMR_0$) | PC2 | 0.134850965 | 0.063716324 | 47.24943859 | 0.141462746 |
| log(Max body depth) -> log($RMR_0$) | PC13 | -2.960409054 | 0.881260253 | 29.76819218 | -0.181449788 |
| log(Max body width) -> log($RMR_0$) | PC14 | -1.517812365 | 0.923816202 | 60.86498126 | -0.084535981 |
| log(Lower jaw length) -> log($RMR_0$) | PC15 | 0.907276433 | 0.44170737 | 48.68498226 | 0.096998637 |
| log(Min caudal peduncle depth) -> log($RMR_0$) | PC16 | -0.384476034 | 0.353786445 | 92.01781476 | -0.048625386 |
| Temperature -> log($RMR_0$) | iPC0 | -0.00187871 | 2.991914 | 159253.6 | -0.00478042 |
| log(Infinite length) -> log($RMR_0$) | iPC1 | 0.10802674 | 2.991914 | 2769.605 | 0.02701592 |
| log($RMR_0$) -> log(Growth coefficient) | PC21 | 0.05837916 | 0.012663124 | 21.69117226 | 0.231203063 |
| log($RMR_0$) -> log(Mortality rate) | PC22 | 0.061831212 | 0.013333797 | 21.56483159 | 0.234909976 |
| log($RMR_0$) -> log(Infinite weight) | PC23 | -0.035634679 | 0.01235603 | 34.67417201 | -0.133254465 |
| log($RMR_0$) -> log(Fecundity) | iPC2 | -0.02133772 | 0.01779804 | 83.41116 | -0.03044145 |
| log($RMR_0$) -> Trophic level | iPC3 | -0.00118572 | 0.0004490948 | 37.87528 | -0.0095525 |

| | | | | | |
|---|---|---|---|---|---|
| log($RMR_0$) -> log(Max age) | iPC4 | -0.04976165 | 0.0146785 | 29.49762 | -0.30697464 |
| log($RMR_0$) -> log(Maturation age invariant) | iPC5 | 0.0218874 | 0.03890677 | 177.7588 | 0.10093606 |
| log($RMR_0$) -> log(Maturation length invariant) | iPC6 | 0.0033883 | 0.006761215 | 199.5459 | 0.03258029 |
| Temperature -> log(K) | PC3 | 0.027907381 | 0.002624492 | 9.404294513 | 0.281229807 |
| Temperature -> log(Mortality) | PC4 | 0.029948825 | 0.003689284 | 12.31862634 | 0.289520964 |
| Temperature -> log(Infinite length) | PC5 | -0.017391136 | 0.002148999 | 12.35686215 | -0.17694811 |
| log(Infinite length) -> log(K) | PC6 | -0.790563285 | 0.024808456 | 3.138073317 | -0.782998508 |
| log(Infinite length) -> log(M) | PC7 | -0.708865467 | 0.034776128 | 4.905885475 | -0.673513141 |
| log(Infinite length) -> log(Body width) | PC8 | -0.00500845 | 0.007164426 | 143.0467618 | -0.022488909 |
| log(Infinite length) -> log(Body depth) | PC9 | -0.01312534 | 0.00788037 | 60.03935842 | -0.053554303 |
| log(Infinite length) -> log(Jaw length) | PC10 | 0.049428991 | 0.012602733 | 25.49664258 | 0.115623016 |
| log(Infinite length) -> log(Peduncle depth) | PC11 | -0.043494823 | 0.014646835 | 33.67489248 | -0.086006709 |
| log(Infinite length) -> log(Infinite weight) | PC12 | 2.943247451 | 0.030706705 | 1.04329337 | 2.752480484 |
| log(K) -> log(Maturity length ratio) | PC24 | 0.163315335 | 0.034918256 | 21.38088049 | 0.396519542 |
| log(K) -> log(Maturity age ratio) | PC25 | -0.432515181 | 0.047325971 | 10.94203687 | -0.503637158 |
| log(M) -> log(Max age) | PC26 | -0.80479829 | 0.018208871 | 2.262538466 | -1.306775685 |
| log(M) -> log(Maturation age ratio) | PC27 | 0.762354026 | 0.04337676 | 5.689844726 | 0.925369425 |
| log(M) -> log(Maturation length ratio) | PC28 | -0.099398255 | 0.034054063 | 34.26022189 | -0.251569755 |
| log(Infinite weight) -> Trophic level | PC29 | 0.033274438 | 0.002579017 | 7.75074453 | 0.071686141 |
| log(Infinite weight) -> log(Fecundity) | PC30 | 0.59879084 | 0.029716305 | 4.962718629 | 0.228446039 |
| H:benthopelagic <-> H:benthopelagic | V[H:benthopelagic ] | 1.222203217 | 1.128684799 | 3.227264521 | |
| H:demersal <-> H:demersal | V[H:demersal] | 1.372694818 | 1.19541574 | 3.179359805 | |
| H:pelagic <-> H:pelagic | V[H:pelagic] | 2.423664029 | 1.591044527 | 5.185602865 | |
| log($RMR_0$) <-> log($RMR_0$) | V[log($RMR_0$)] | 2.202400861 | 2.004507501 | 3.414330025 | |
| log(Maturation age ratio) <-> log(maturation age ratio) | V[log(maturation age ratio)] | 0.103560006 | 0.337615419 | 3.296830734 | |
| log(Maturation length ratio) <-> log(Maturation length ratio) | V[log(maturation length ratio)] | 0.023820432 | 0.165993095 | 4.518422218 | |
| log(maximum age)<-> log(Maximum age) | V[log(maximum age)] | 0.057873662 | 0.24915902 | 2.748643276 | |
| log(K) <-> log(K) | V[K] | 0.140418679 | 0.48201672 | 2.171687642 | |
| log(Infinite weight) <-> log(Infinite weight) | V[log(Infinite weight)] | 0.157499263 | 0.491787037 | 2.225163415 | |
| log(Mortality) <-> log(Mortality) | V[Mortality] | 0.152583988 | 0.518486938 | 2.744396865 | |
| Trophic level <-> Trophic level | V[Trophic level] | 0.033933571 | 0.192724522 | 0.607253487 | |
| log(Infinite length) <-> log(Infinite length) | V[log(Infinite length)] | 0.137744247 | 0.382020452 | 1.760437813 | |
| log(Fecundity) <-> log(Fecundity) | V[log(Fecundity)] | 1.082085453 | 1.091290771 | 2.383597294 | |
| log(Peduncle depth) <-> log(Peduncle depth) | V[log(Peduncle depth)] | 0.035227673 | 0.272606214 | 1.214195093 | |
| log(Jawlength) <-> log(Jawlength) | V[log(Jawlength)] | 0.025173777 | 0.230088555 | 1.227043476 | |
| log(Body depth) <-> log(Body depth) | V[log(Body depth)] | 0.008273816 | 0.131989026 | 1.203815908 | |
| log(Body width) <-> log(Body width) | V[log(Body width)] | 0.00683193 | 0.120466549 | 1.196036896 | |
| Temperature <-> Temperature | V[Temperature] | 14.25966783 | 3.846206967 | 0.643608906 | |

236

### S3 - Supplementary material 3 - Cross-Validation (CV)

10 folds cross-validation percentage variance explained equation (Thorson et al., 2023), where $y_{i,t}$ corresponds to the observed trait $t$ value of the species $i$, and $\bar{y}_t$ to the mean value of the trait. $\beta_i^{(t)}$ is the predicted trait values of the species $i$ for trait $t$, with the PSEM method.

$$PVE_t = 1 - \frac{\Sigma_{i=1}^{n_i}\left(y_{i,t} - \beta_i^{(t)}\right)^2}{\Sigma_{i=1}^{n_i}\left(y_{i,t} - \underline{y_t}\right)^2} \qquad (3)$$

Jackknife cross-validation standard error estimate, with $\hat{\theta}_{(i)}$ the parameter of interest, inferred without the i^th level of $RMR_0$, and $\hat{\theta}_{(.)}$the mean of the $n\hat{\theta}_{(i)}$. The relative standard error was estimated as the standard error divided by the inferred trait value with the complete dataset (McIntosh (2016)) :

$$SE(\hat{\theta})_{jack} = \sqrt{\frac{n-1}{n}\Sigma_{i=1}^{n} \left(\hat{\theta}_{(i)} - \hat{\theta}_{(.)}\right)^2} \qquad (4)$$

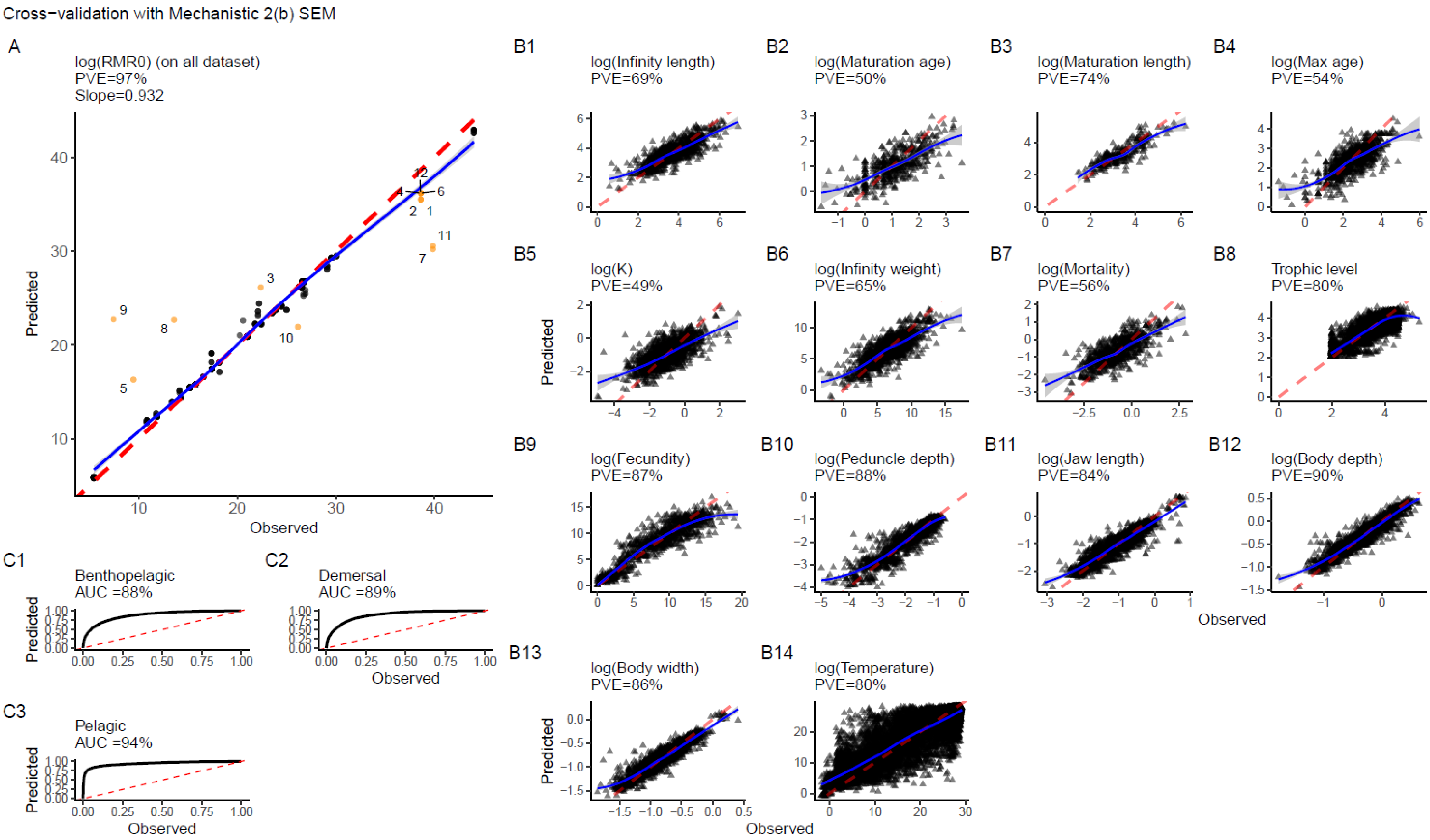

Cross−validation with Mechanistic 2(b) SEM
A
log(RMR0) (on all dataset)
PVE=97%
Slope=0.932
Predicted
Observed
B1
log(Infinity length)
PVE=69%
B2
log(Maturation age)
PVE=50%
B3
log(Maturation length)
PVE=74%
B4
log(Max age)
PVE=54%
B5
log(K)
PVE=49%
B6
log(Infinity weight)
PVE=65%
B7
log(Mortality)
PVE=56%
B8
Trophic level
PVE=80%
B9
log(Fecundity)
PVE=87%
B10
log(Peduncle depth)
PVE=88%
B11
log(Jaw length)
PVE=84%
B12
log(Body depth)
PVE=90%
B13
log(Body width)
PVE=86%
B14
log(Temperature)
PVE=80%
C1
Benthopelagic
AUC =88%
C2
Demersal
AUC =89%
C3
Pelagic
AUC =94%


249

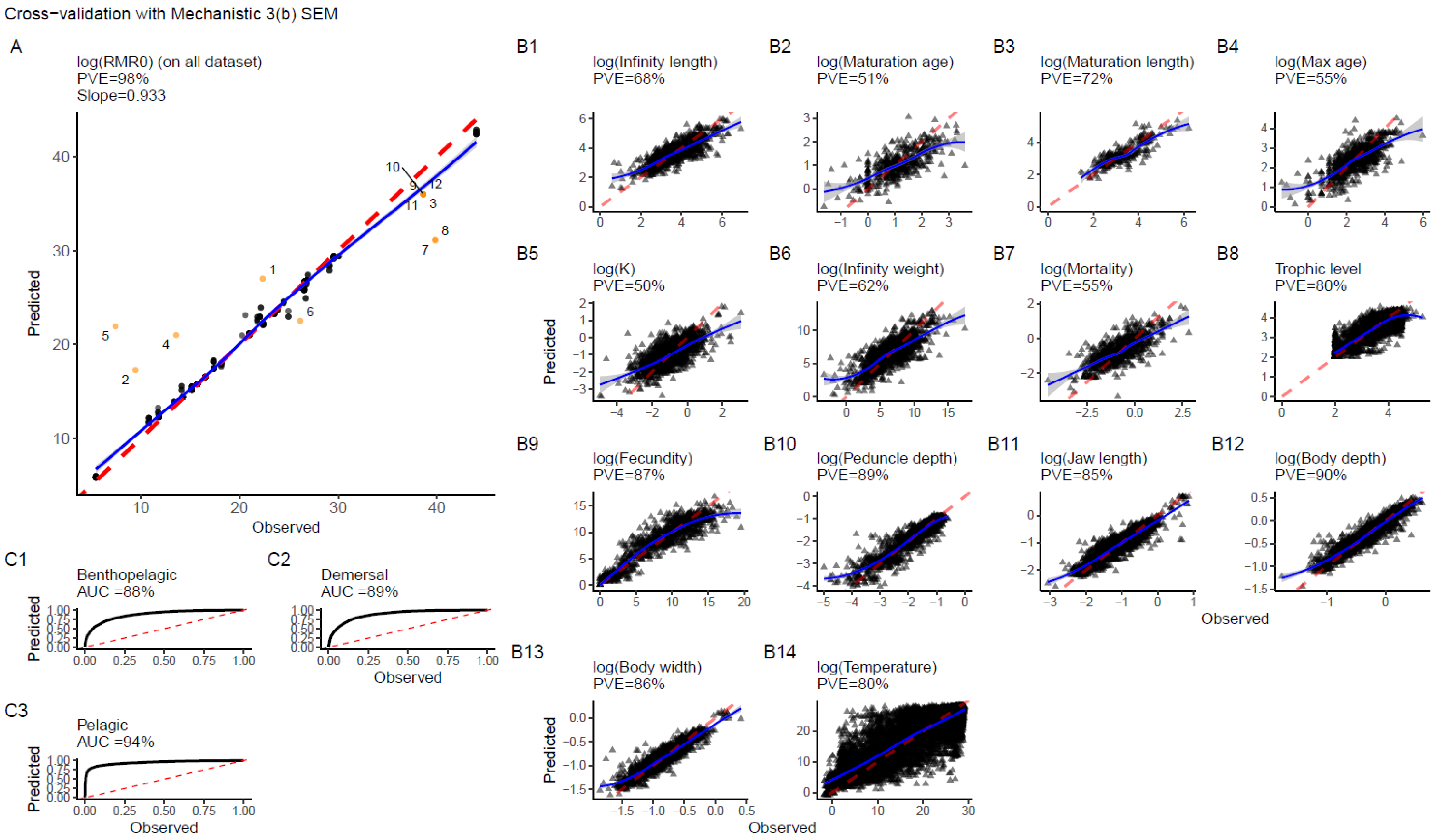

Cross−validation with Mechanistic 3(b) SEM
A
log(RMR0) (on all dataset)
PVE=98%
Slope=0.933
Predicted
Observed
B1
log(Infinity length)
PVE=68%
B2
log(Maturation age)
PVE=51%
B3
log(Maturation length)
PVE=72%
B4
log(Max age)
PVE=55%
B5
log(K)
PVE=50%
B6
log(Infinity weight)
PVE=62%
B7
log(Mortality)
PVE=55%
B8
Trophic level
PVE=80%
B9
log(Fecundity)
PVE=87%
B10
log(Peduncle depth)
PVE=89%
B11
log(Jaw length)
PVE=85%
B12
log(Body depth)
PVE=90%
B13
log(Body width)
PVE=86%
B14
log(Temperature)
PVE=80%
C1
Benthopelagic
AUC =88%
C2
Demersal
AUC =89%
C3
Pelagic
AUC =94%

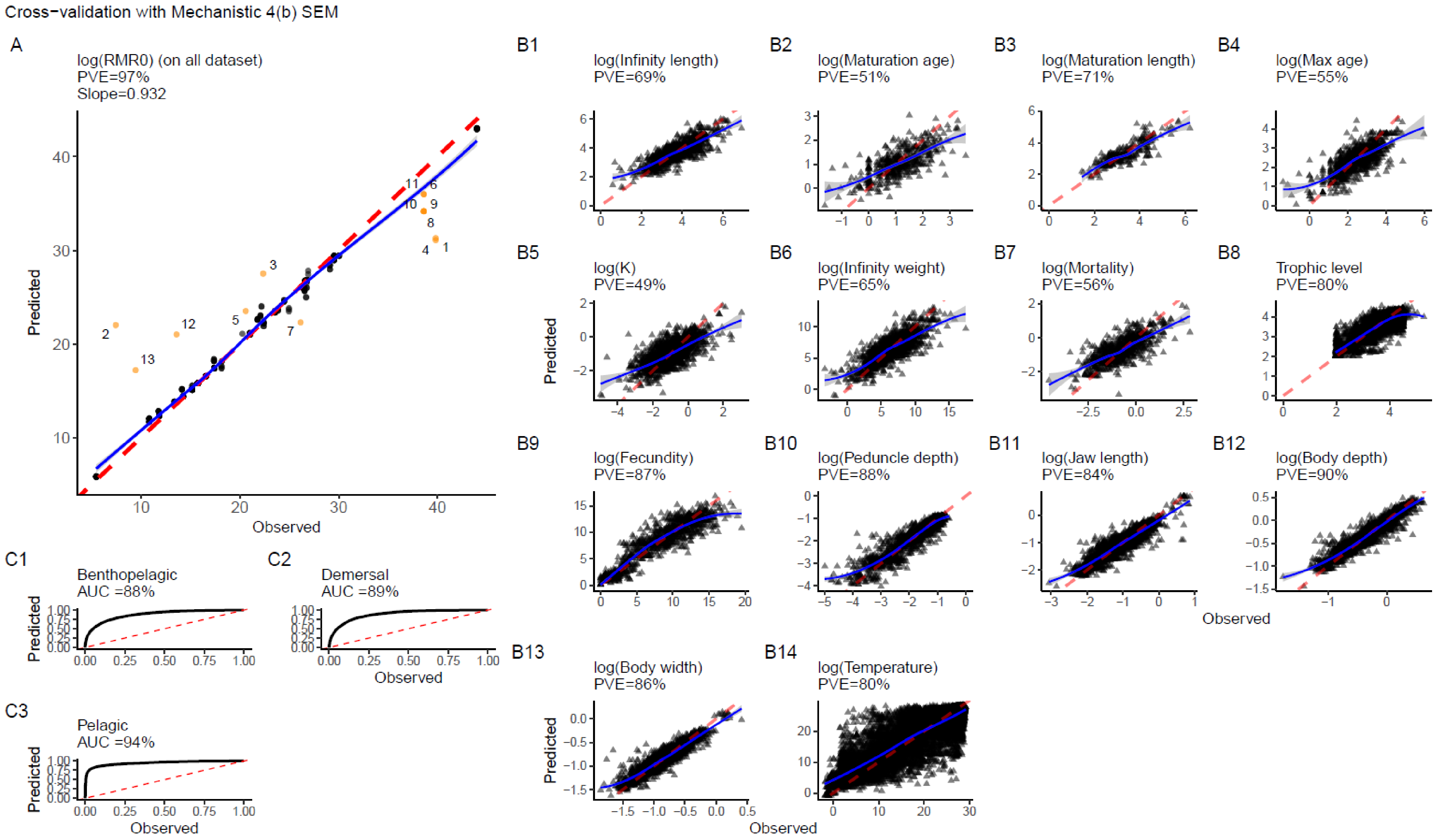
Cross−validation with Mechanistic 4(b) SEM
A
log(RMR0) (on all dataset)
PVE=97%
Slope=0.932
Predicted
Observed
B1
log(Infinity length)
PVE=69%
B2
log(Maturation age)
PVE=51%
B3
log(Maturation length)
PVE=71%
B4
log(Max age)
PVE=55%
B5
log(K)
PVE=49%
B6
log(Infinity weight)
PVE=65%
B7
log(Mortality)
PVE=56%
B8
Trophic level
PVE=80%
B9
log(Fecundity)
PVE=87%
B10
log(Peduncle depth)
PVE=88%
B11
log(Jaw length)
PVE=84%
B12
log(Body depth)
PVE=90%
B13
log(Body width)
PVE=86%
B14
log(Temperature)
PVE=80%
C1
Benthopelagic
AUC =88%
C2
Demersal
AUC =89%
C3
Pelagic
AUC =94%
Predicted
Observed

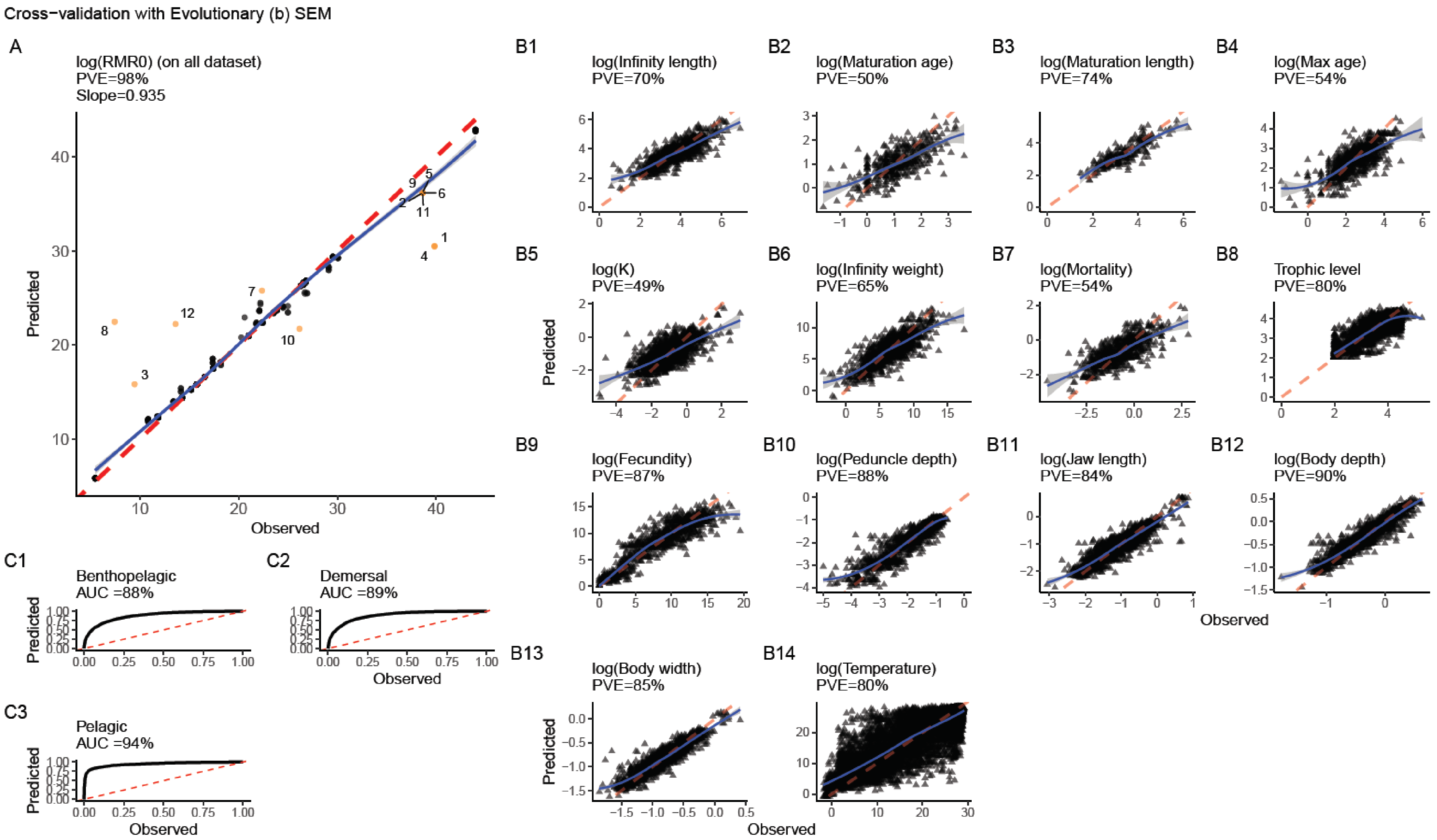
Cross−validation with Evolutionary (b) SEM
A
log(RMR0) (on all dataset)
PVE=98%
Slope=0.935
Predicted
Observed
B1
log(Infinity length)
PVE=70%
B2
log(Maturation age)
PVE=50%
B3
log(Maturation length)
PVE=74%
B4
log(Max age)
PVE=54%
B5
log(K)
PVE=49%
B6
log(Infinity weight)
PVE=65%
B7
log(Mortality)
PVE=54%
B8
Trophic level
PVE=80%
B9
log(Fecundity)
PVE=87%
B10
log(Peduncle depth)
PVE=88%
B11
log(Jaw length)
PVE=84%
B12
log(Body depth)
PVE=90%
B13
log(Body width)
PVE=85%
B14
log(Temperature)
PVE=80%
C1
Benthopelagic
AUC =88%
C2
Demersal
AUC =89%
C3
Pelagic
AUC =94%

Figure S3.1 | Four ten-fold cross-validation output plots comparing inferred trait values with observed values over the complete constructed dataset. Each plot refers to a different SEM (b) representing the SEM where all morphometric traits explain trophic level. The red dotted line indicates the 1:1 relationship, and the blue line represents a Loess regression curve. Each point corresponds to a species. Quantitative and qualitative traits are plotted separately. A: $RMR_0$ cross-validation plot. Orange dots labeled as outliers have a Cook's distance greater than four times the average. B: Qualitative traits cross-validation plots: habitat traits. The plots represent the performance of the binary trait discrimination (1 and 0). The AUC indicator reflects this performance: the closer to 100%, the better the discrimination power of the trait value. C: Quantitative traits cross-validation plots. The better the inference, the closer the data points to the red-dotted curve. The percentage variance explained (PVE) is the sum of the squared differences between observed and inferred trait values divided by the sum of the squared differences between observed and the mean of the observed trait values (S3-Eq vi).

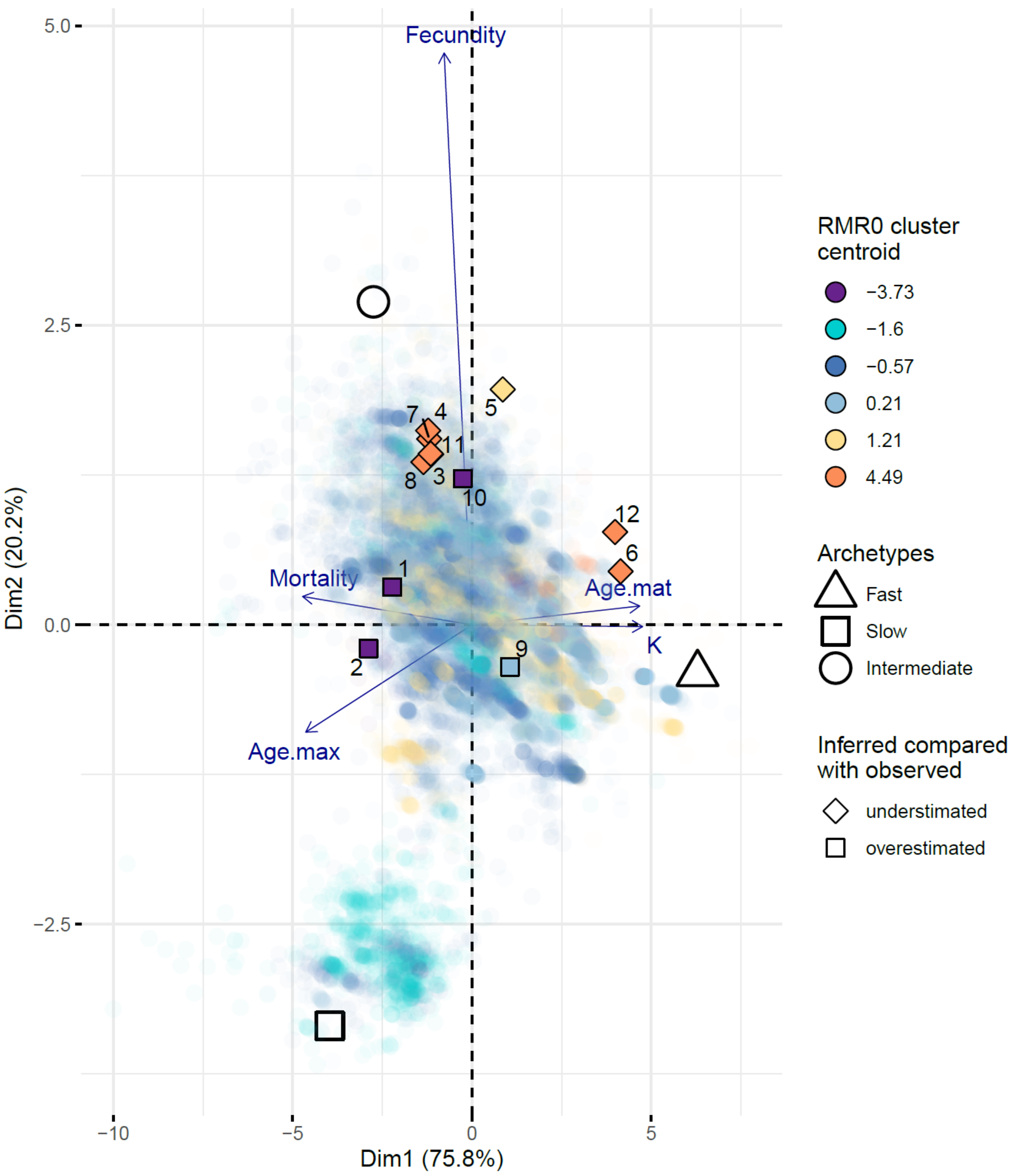


Figure S3.2 | Across-traits cross-validation outliers, plotted in the first and second dimensions of a PCA. The PCA includes the following traits: maximum age (Age.max), maturation age (Age.mat), growth coefficient (K) and mortality rate (Mortality). Triangle, square and circle shapes indicate archetypes corresponding to fast, slow, and intermediate life-history strategies, classified based on the growth coefficient: fast = high K, slow = low K and intermediate = intermediate K. Each colored point represents a species associated with a standardized $RMR_0$ cluster, determined using the k-means algorithm (k=6) (r-package *Ckmeans.1d.dp* v4.3.5, Wang et Song, 2011). Points in the background show the 18,214 species with observed or inferred functional trait values. Black-circled reversed triangles and diamonds indicate species with overestimated or underestimated $RMR_0$ outliers. Outliers are defined as species with Cook's distance greater than four times the mean Cook's distance. Point numbers correspond to the following species: (1) Channa brunnea, (2) Chaenocephalus aceratus, (3) Pseudopleuronectes obscurus, (4) Pseudopleuronectes yokohamae, (5) Chanos chanos, (6) Gillichthys seta, (7) Pseudopleuronectes americanus, (8) Pseudopleuronectes herzensteini, (9) Gilchristella aestuaria, (10) Mugil capurrii, (11) Pseudopleuronectes schrenki, (12) Gillichthys mirabilis.

| | Q10 | Q25 | Median | Q75 | Q90 |
|---|---|---|---|---|---|

| | | | | | |
|---|---|---|---|---|---|
| Infinite length | 0.01446876 | 0.03302718 | 0.07844345 | 0.15565941 | 0.28698445 |
| Infinite weight | 0.02138109 | 0.0886544 | 0.20254794 | 0.4053247 | 0.82560658 |
| Maturation length | 0.14116105 | 0.22343173 | 0.34150977 | 0.54629849 | 0.85729974 |
| Maturation age | 0.87744683 | 1.88921476 | 4.06346729 | 9.09602155 | 20.7557463 |
| Maximum age | 0.29295661 | 0.64663593 | 1.24539213 | 2.29881831 | 4.15101404 |
| Fecundity | 0.00772199 | 0.02632265 | 0.06185358 | 0.12025974 | 0.21471118 |
| K | 0 | 0.78638815 | 1.97235807 | 4.68870683 | 12.937439 |
| Mortality | 0.98026085 | 2.15603049 | 4.51426711 | 9.95510319 | 25.0981742 |
| Max body width | 0 | 0.00324542 | 0.01465691 | 0.04647677 | 0.33143092 |
| Max body depth | 0 | 0.00962466 | 0.0561822 | 0.23465751 | 1.80905886 |
| Jaw length | 0 | 0.00470005 | 0.02104149 | 0.06891973 | 0.57526606 |
| Peduncle depth | 0 | 0.0024029 | 0.01142911 | 0.0356575 | 0.28685808 |
| Temperature | 0 | 0 | 0 | 0.00097144 | 0.00587918 |
| $RMR_0$ | 7.66359358 | 9.07135496 | 11.5007839 | 16.7587697 | 45.942652 |

Table S3.1 | Quantiles (Q) of jackknife relative standard error (RSE) estimates. The jackknife assessed the effect of removing one of the 43 genus-level observed $RMR_0$ values on the PSEM. Each sub-sample dataset corresponds to the functional traits dataset prior to inference, excluding the species-level data of the selected genus. RSEs were calculated for each trait and each species across the 43 inferred datasets. No quantiles were estimated for traits without inference (habitat and trophic level) because these datasets were complete.

### S4 - Supplementary material 4 - Archetypal analysis (AA) and $RMR_0$ gradient

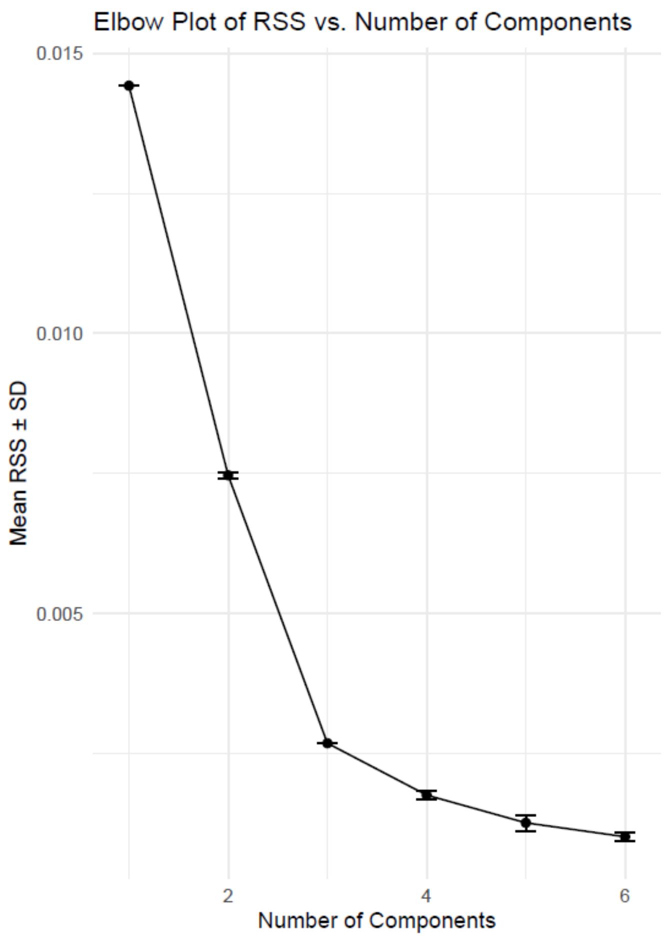


Figure S4.1 | Elbow plot from 30 archetypal analysis replicates, considering one to six clusters. The fit of the archetypes improves as the residual sum of squares (RSS) decreases, a criterion that balances model complexity and goodness of fit. The elbow is located at the third archetype: three archetypes are sufficient to represent the dataset.

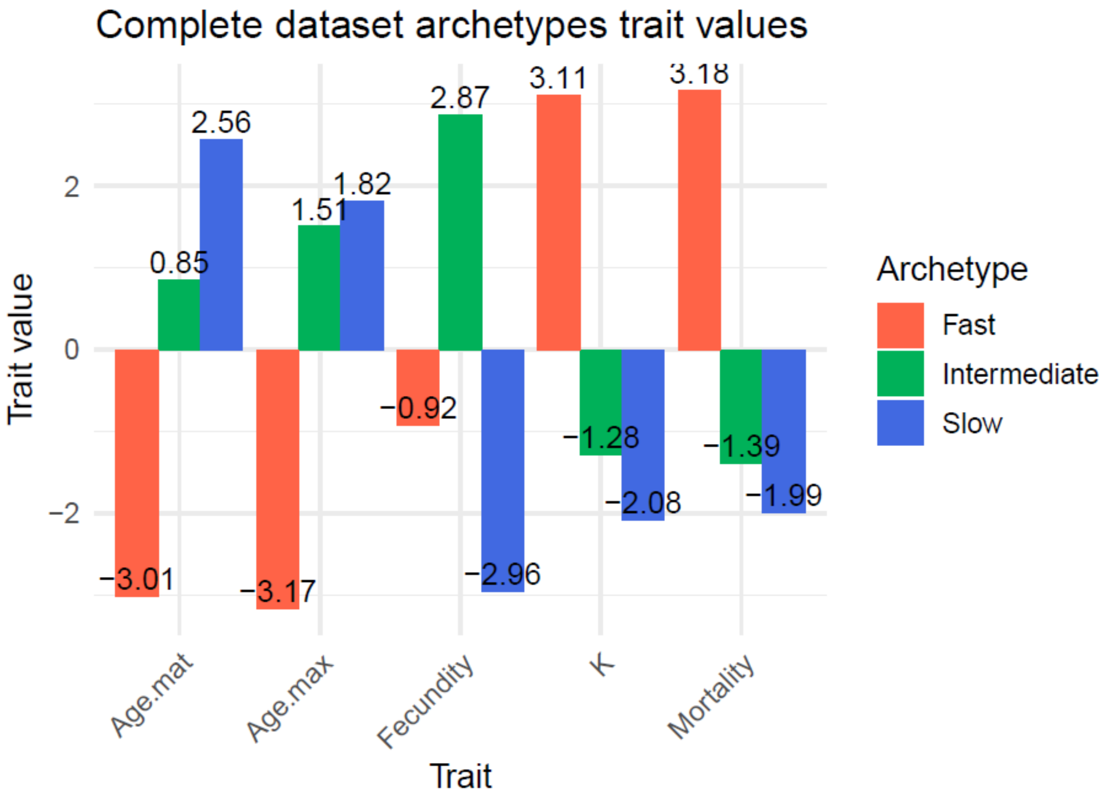


Figure S4.2 | Functional trait groups from the archetypal analysis, represented by their trait values. The color of the bars indicates the archetype group along the fast-slow continuum. Archetype are identified based on the growth coefficients: highest = fast archetype, lowest = slow archetype and intermediate = intermediate archetype.

Table S4.1 | Linear regression of $RMR_0$ against PC1, PC2 and their interaction. Estimates correspond to the partial regression coefficients, interpreted as standardized partial regression coefficients since the PCA was performed on standardized trait values. Standard errors and p-values (F-tests based on sum-of-squares) are provided. Relative importance is obtained as the product between the partial regression coefficient and the square root of the eigenvalue of the corresponding PC (or the product of the eigenvalues for the interaction) normalized by the sum of all products. This metric accounts for both the strength of the relationship (partial regression coefficient) and the variance explained (eigenvalue) in the influence of each PC on predicted variability in $RMR_0$.

| | **Estimate** | **Standard error** | **p-value** | **Relative importance** |
|---|---|---|---|---|
| PCA1 | 0.13 | 1e-03 | <2e-16 | 46% |
| PCA2 | 0.29 | 0.01 | <2e-16 | 53% |
| PCA1:PCA2 | 0.01 | 1e-03 | 0.013 | 1% |
